# Visualizing translation dynamics at atomic detail inside a bacterial cell

**DOI:** 10.1101/2021.12.18.473270

**Authors:** Liang Xue, Swantje Lenz, Maria Zimmermann-Kogadeeva, Dimitry Tegunov, Patrick Cramer, Peer Bork, Juri Rappsilber, Julia Mahamid

## Abstract

Translation is the fundamental process of protein synthesis and is catalysed by the ribosome in all living cells. Here, we use cryo-electron tomography and sub-tomogram analysis to visualize the dynamics of translation inside the prokaryote *Mycoplasma pneumoniae*. We first obtain an in-cell atomic model for the *M. pneumoniae* ribosome that reveals distinct extensions of ribosomal proteins. Classification then resolves thirteen ribosome states that differ in conformation and composition and reflect intermediates during translation. Based on these states, we animate translation elongation and demonstrate how antibiotics reshape the translation landscape inside cells. During translation elongation, ribosomes often arrange in a defined manner to form polysomes. By mapping the intracellular three-dimensional organization of translating ribosomes, we show that their association into polysomes exerts a local coordination mechanism that is mediated by the ribosomal protein L9. Our work demonstrates the feasibility of visualizing molecular processes at atomic detail inside cells.

---

Translation of genetic information via messenger RNAs (mRNA) into proteins is carried out by the ribosome, one of the most primordial macromolecular machines in cells^1,2^. The process of translation has been subject to extensive structural studies since the first observation of ribosomes under an electron microscope^3-9^. The ribosome consists of a small and a large subunit (30S and 50S in prokaryotes), which form the aminoacyl (A), peptidyl (P) and exit (E) transfer RNA (tRNA) binding sites at their interface. The translation process can be divided into four phases; initiation, elongation, termination, and recycling. During the elongation phase, the ribosome undergoes a fundamental cycle to add one amino acid to the nascent peptide chain. Each fundamental elongation cycle can be subdivided into three steps: decoding, peptidyl transfer and translocation. These steps involve structural changes in the ribosome that include subunit rotations, elongation factor association and tRNA accommodation^6-8^. Many intermediates during the elongation cycle have been identified based on structures of ribosomes that were isolated from model bacteria such as *Escherichia coli* and *Thermus thermophilus*, and were often trapped in different functional states by antibiotics, GTP analogues or mutations^7-10^. Although ribosomes have been visualized inside cells^11-15^, a detailed investigation of the translation process within the native cellular context is lacking. Here, we harnessed recent advances in cryo-electron tomography (cryo-ET) and sub-tomogram analysis^16,17^ to visualize parts of the translation process at atomic detail inside *Mycoplasma pneumoniae*, a prokaryotic minimal cell model^11^.

## The in-cell *M. pneumoniae* ribosome structure reveals distinct protein extensions

To investigate the structural details of the translation machinery, we first obtained 3.5 Å ribosome maps from *M. pneumoniae* cellular tomograms (Fig. 1a-b, Extended Data Fig. 1, Supplementary Discussion). Focused refinements on the 30S and 50S subunits improved the densities and revealed well-resolved ribosomal RNA bases and amino acid side chains (Fig. 1b, Extended Data Fig. 2a-c). This allowed us to build an atomic model for the *M. pneumoniae* ribosome, which shows high structural similarity to other bacterial ribosomes, but also reveals several new features (Fig. 1c, Extended Data Fig. 2d-g). Specifically, eleven of the 52 ribosomal proteins have extended sequences compared to *E. coli* (Extended Data Fig. 3a, Supplementary Discussion). Most of these extensions are predicted to be disordered, but those of the ribosomal proteins S6, L22 and L29 form secondary structures and were built in our atomic model (Extended Data Fig. 2d-e, 3a). We found such extensions to be common throughout the bacterial kingdom (Extended Data Fig. 3b, Supplementary Discussion), possibly representing ribosome diversity in adaptation to different environments and lifestyles^1,18^. Although putative functions of these extensions remain largely elusive, a transposon mutation screen in *M. pneumoniae* shows that their disruption affects cellular fitness or survival (Supplementary Discussion)^19^. Our structure provides the basis to investigate the conformational and compositional changes of ribosomes during translation in *M. pneumoniae* cells.

**Fig. 1.**
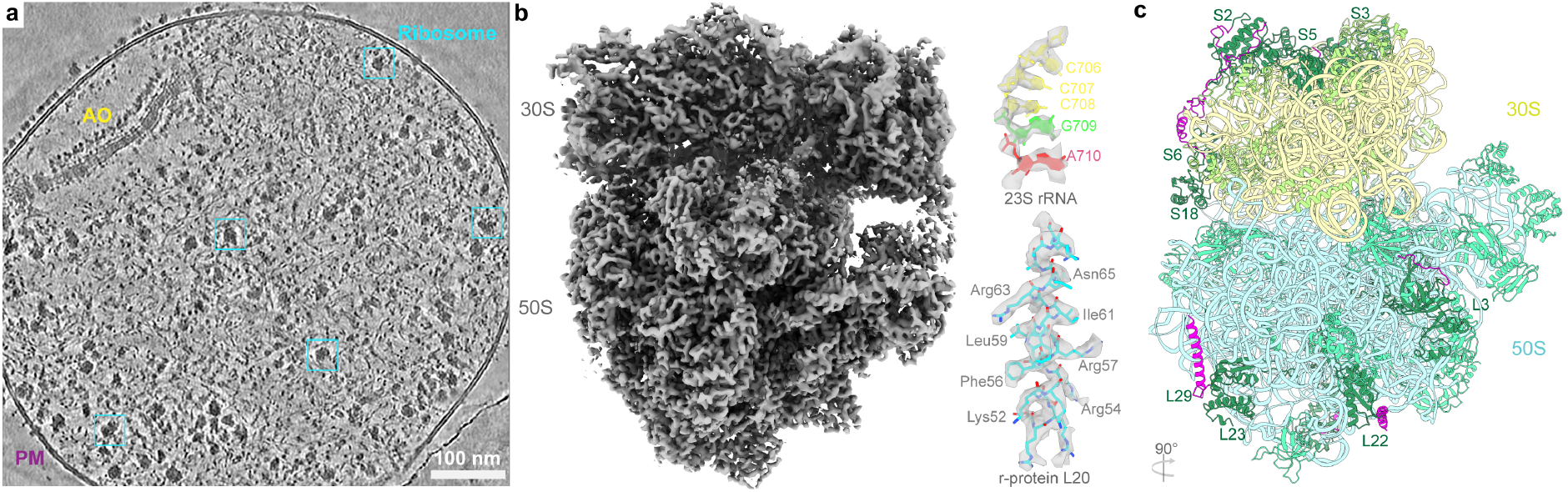
Ribosome structure in *M. pneumoniae* cells. **a**, A denoised tomographic slice of a *M. pneumoniae* cell. AO: attachment organelle; PM, plasma membrane. Examples of ribosomes are highlighted in cyan boxes. **b**, 3.5Å in-cell ribosome map (left) shows well-resolved RNA bases and amino acid side chains (right). **c**, Atomic model of the *M. pneumoniae* ribosome shows structural similarity to other bacterial ribosomes. Eleven ribosomal proteins (dark green) have sequence extensions (magenta).

## Structural dynamics of translation elongation inside cells

To dissect the translation process inside cells, we performed classification^20^ of 101,696 ribosomes from 356 cellular tomograms that resulted in 13 distinct structures (Extended Data Fig. 4, Methods). Ten classes determined at resolutions ranging from 5 to 10 Å were assigned to the elongation phase based on elongation factor and tRNAs (“EF + tRNAs’’) binding to the ribosome (Fig. 2a, Extended Data Figs. 4-7). The remaining three classes represent 70S with single P/E-site tRNA, 50S in complex with the ribosome recycling factor (RRF), and free 50S subunits (Supplementary Discussion). The ten classes within the elongation phase account for 70% of all localized ribosomes, consistent with the expectation that the majority of 70S ribosomes are engaged in the elongation phase, which lasts considerably longer than the initiation, termination and recycling phases^8,21,22^.

**Fig. 2.**
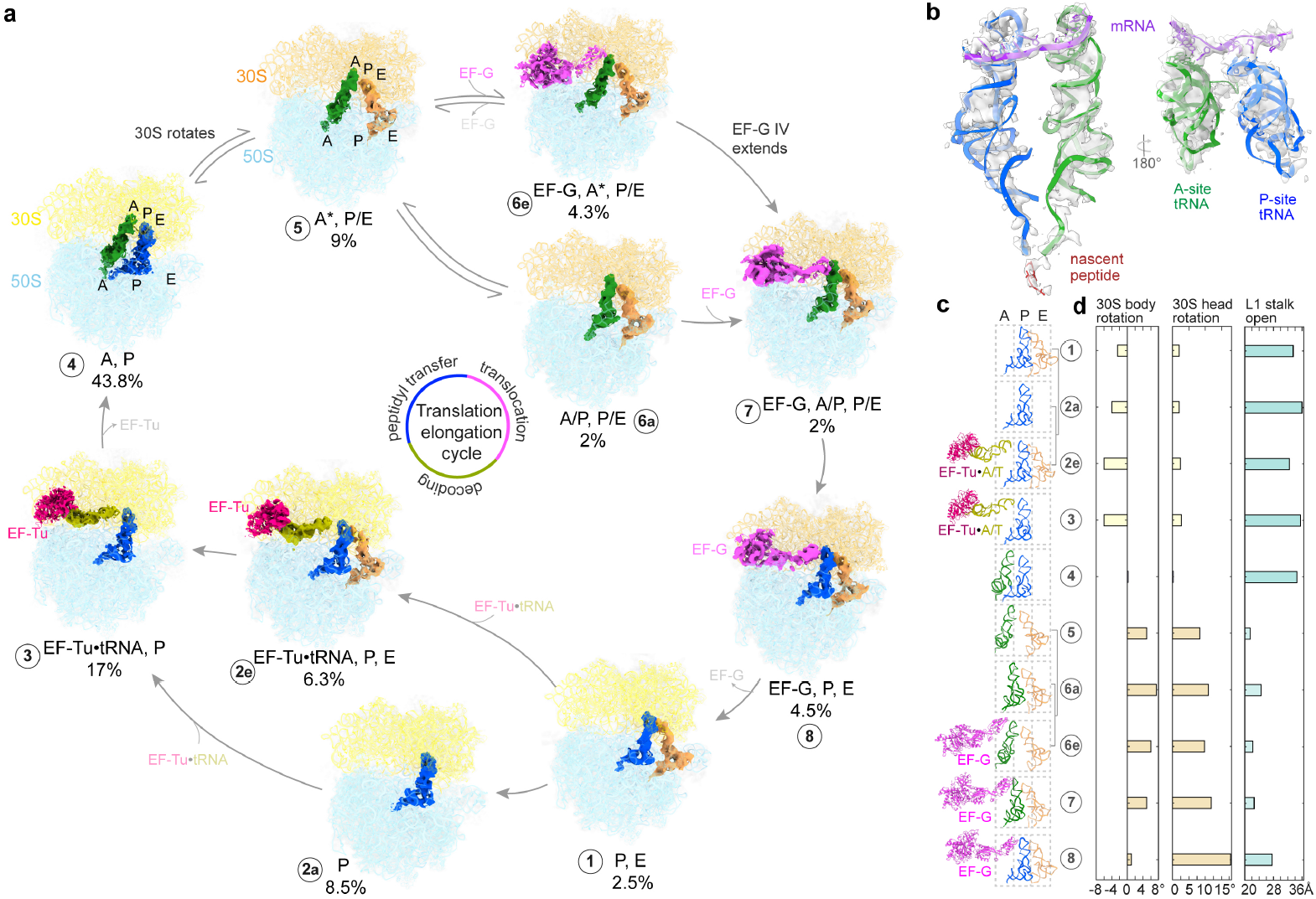
Ribosome classification reconstructs the translation elongation cycle. **a**, Ten 70S structures determined inside *M. pneumoniae* cells represent translation elongation intermediate states, characterized by different elongation factor and tRNA binding. The occurrence frequencies for different intermediates are calculated from the classified particle numbers. **b**, Densities for mRNA, tRNAs and the nascent peptide chain are well-resolved in the most populated “A, P” state. **c**, Trajectories of elongation factors EF-Tu, EF-G, and the A-P-E site tRNAs along translation elongation. **d**, Major conformational changes of the ribosome along the elongation trajectory: 30S body rotation, 30S head rotation (in degrees) and L1 stalk opening.

The identified elongation-related classes can be ordered to reconstruct the translation elongation cycle^7,8,23^. By flexible fitting of atomic models, we could delineate conformational and compositional changes in the ribosome complex, including rotations of 30S body and head, movement of tRNAs through the A-P-E sites, dynamics of the L1 stalk, and coordination of elongation factors (Fig. 2, Extended Data Fig. 7a-j, Movie 1, 2). The number of particles within each of the structural classes also provides the relative abundance of intermediates that reflects a steady state distribution determined by their relative rates of formation and depletion inside the cells (Fig. 2a, Extended Data Fig. 4). The pre-translocational “A, P” class was the most populated and was determined at the highest resolution, showing clear density for mRNA, tRNAs and the nascent peptide (Fig. 2b). In the following states, we observed oscillation of A- and P-site tRNAs into the hybrid A/P and P/E state coupled with 30S subunit rotations (Fig. 2a, c, d, Extended Data Fig. 7a-j), as previously reported^3,10,23-27^. We also determined a partial hybrid “A*, P/E” state (Fig. 2a, d), where only the A-site tRNA marginally moves^25,26^. Interestingly, EF-G was found to bind to the ribosome in either the partial hybrid or the full hybrid states. In the partial hybrid state, EF-G is less extended and its domain IV does not overlap with the A site compared to the full hybrid state, possibly representing an intermediate transiting to the fully elongated conformation^10,23,28,29^. Two EF-Tu-associated structures were also determined, with and without E-site tRNA (Fig. 2a, Extended Data Fig. 7c, d). This suggests that binding of EF-Tu•tRNA to the ribosome and disassociation of E-site tRNA are uncoupled. The “P, E” class has a relatively low abundance, suggesting that the E-site tRNA is not stable and tends to disassociate quickly after translocation^8,30^. This is in agreement with previous single-molecule fluorescence studies showing rapid release of the E-site tRNA^31,32^, and helps resolving a long-standing controversy over its disassociation time point^7,33,34^. In summary, these results recapitulate major steps in translation elongation that have been defined by *in vitro* studies and reconstruct the structural dynamics of the elongation cycles inside cells. Furthermore, with these data, a probability map for the occurrence of intermediate states can be derived to illustrate a putative energy landscape of the translation elongation cycle in cells. Energy landscapes of translation are suggested to be rugged^25-27,34-37^; the thermodynamics and kinetics of translation have been shown to vary depending on factors such as Mg^2+^ concentrations^38,39^, EF-G^30,35^ and temperature^26,36^. In *M. pneumoniae*, the intracellular concentrations of ribosomes, accessory factors and substrates can be up to ten times lower than those in *E. coli* (Supplementary Discussion). Therefore, it is possible that energy landscapes of translation vary for different organisms, cell types and conditions. Applying the analysis introduced here to other cell types and conditions can deepen our understanding of the translation process within the native cellular context.

## Antibiotics alter translation landscapes in cells

The ribosome is one of the most important targets for antibiotics, many of which are known to stall translation by stabilizing certain intermediates in the process^40^. To investigate the effects of antibiotics on the cellular translation machinery, we analysed ribosomes in cells that were shortly (15-20 minutes) treated with two representative ribosome-targeting antibiotics: chloramphenicol (Cm) that binds to the peptidyl transfer center in the 50S subunit and inhibits peptide bond formation^41-43^, and spectinomycin (Spt) that binds to the 30S neck and blocks translocation^44-47^. Overall, our sub-tomogram analysis of the antibiotic-treated cells resulted in 17 ribosome maps mostly resolved at 4 to 10 Å resolutions (Extended Data Fig. 8-10), and shows that translation landscapes are dramatically reshaped under the different antibiotic treatments (Fig. 3a, Supplementary Discussion).

**Fig. 3.**
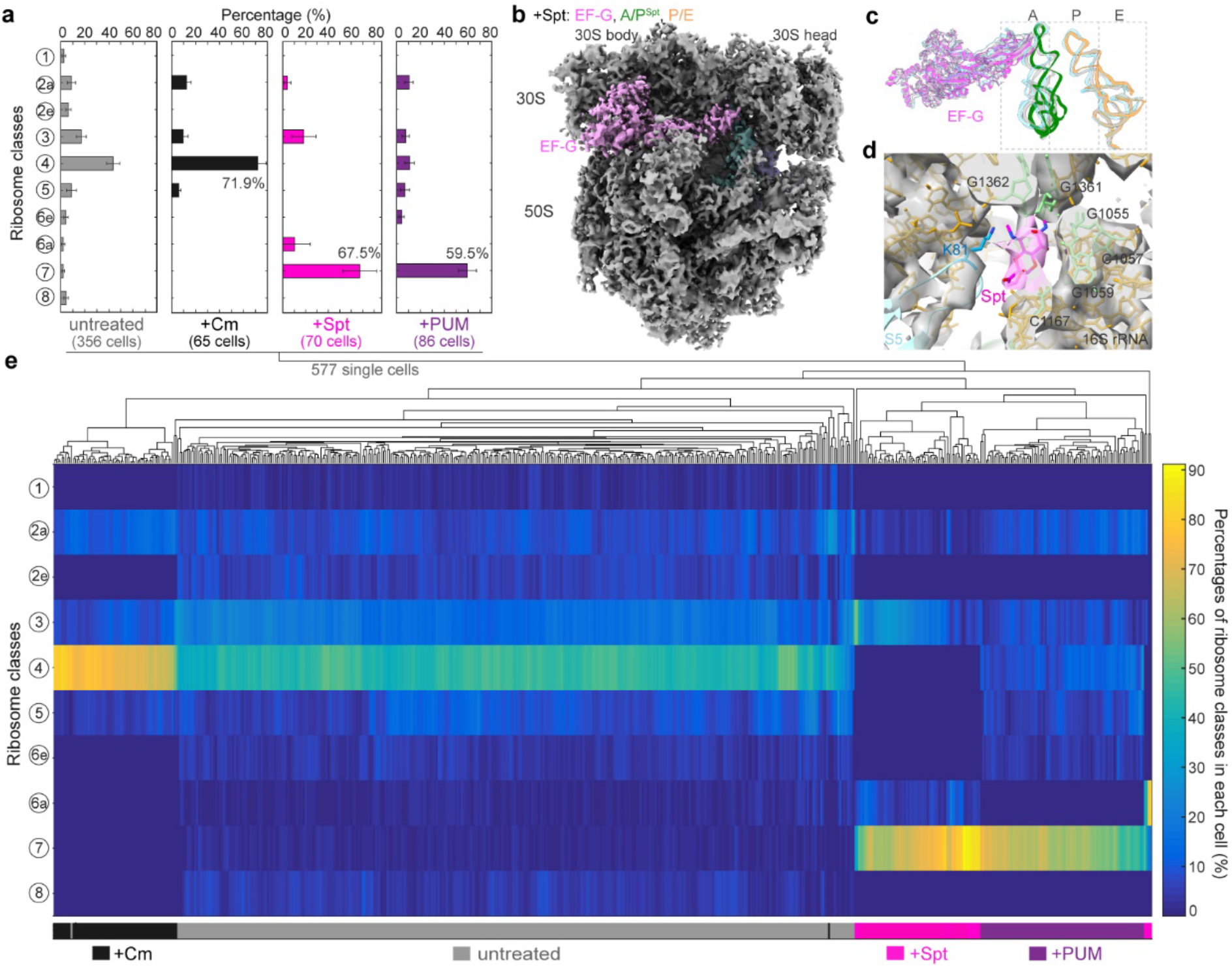
Antibiotics induce distinct translation elongation landscapes in cells. **a**, Distribution of translation elongation intermediates in untreated and three antibiotic-treated cells. +Cm, chloramphenicol-treated; +Spt, spectinomycin-treated; +PUM, pseudouridimycin-treated. **b**, Ribosomes in Spt-treated cells are largely stalled in “EF-G, A/P^Spt^, P/E’’ state. **c**, The major state in Spt-treated cells is similar to the “EF-G, A/P^PUM^, P/E’’ class in PUM-treated cells (light blue) and the “EF-G, A/P, P/E’’ class in untreated cells (light grey). **d**, The Spt molecule (magenta) is well-resolved and built in the “EF-G, A/P^Spt^, P/E’’ ribosome model. It is surrounded by several 16S rRNA bases and the loop 2 of ribosomal protein S5 near the 30S neck. **e**, Single cell clustering analysis based on translation elongation states of 577 cells under native conditions and different antibiotic treatments.

In Cm-treated cells, 72% of 70S ribosomes are trapped in the expected “A, P” state, owing to the inhibition of peptidyl transfer^40^ (Fig. 3a-d, Extended Data Fig. 8). The Cm molecule was resolved in its canonical binding site^41-43^ (Extended Data Fig. 8d, g). Interestingly, 28% of ribosomes were found in several other elongation states, either prior to or following the major “A, P” state (Fig. 3a, Extended Data Fig. 8). No significant class of free 50S was detected, possibly owing to the inhibition of ribosome dissociation mediated by EF-G and RRF by Cm^48^.

Spt is suggested to inhibit translocation by blocking head swivel of the 30S subunit^44-47^. We show that 67.5% of 70S ribosomes are stalled in the “EF-G, A/P^Spt^, P/E” state, a state similar to the pre-translocational “EF-G, A/P, P/E” state in untreated cells (Fig. 3a-d, Extended Data Fig. 9). The Spt molecule density is clearly visible near helix 34 within the 30S neck region (Fig. 3d, Extended Data Fig. 9d, g), consistent with its reported binding site^44,45^. These results ratify that Spt inhibits translocation by impeding dynamics of the 30S subunit. Similar to Cm, ribosomes in three additional elongation states could be detected with lower frequencies (Fig. 3a, Extended Data Fig. 9).

Interestingly, perturbation of other molecular pathways also affects translation in cells. RNA polymerase stalled by pseudouridimycin (PUM) can physically block mRNA translocation in the ribosome that collides with it during transcription-translation coupling^16,49^. Consistently, 59.5% of 70S ribosomes in PUM-treated cells were found in the “EF-G, A/P^PUM^, P/E” state (Fig. 3a, Extended Data Fig. 10), which resembles the pre-translocational “EF-G, A/P, P/E” state in untreated cells and the “EF-G, A/P^Spt^, P/E” state in Spt-treated cells (Fig. 3c). The finding that physically obstructing mRNA translocation by PUM-stalled RNA polymerase and chemically impeding 30S head dynamics by Spt lead to similar structures further confirms that mRNA translocation and 30S rotations are directly coupled.

The observation that antibiotics treatment also resulted in minor fractions of states not expected based on their specific binding prompted us to investigate possible cell-to-cell variability in response to the antibiotics. To this end, we performed clustering analysis of single cells based on translation elongation profiles for 577 tomograms of individual cells under native and antibiotic treatment conditions. The analysis resulted in four major clusters in accordance with the four treatment groups, demonstrating small cell-to-cell variability within each cluster (Fig. 3e). In summary, our results show that the translation landscape in cells is reshaped by small molecules specifically binding to ribosomes, as well as to other targets. The presence of minor elongation states under antibiotic treatment is reminiscent of recent works showing ongoing slow translation in antibiotic-treated cells and context-dependent inhibition^50-52^. For example, Cm inhibition is affected by specific residues of the nascent peptide^50,51^. Most antibiotics, including Cm and Spt, inhibit cell growth but do not immediately kill the cell^53^. It is possible that the reshaped translation landscapes by antibiotics lead to imbalance in protein synthesis, which in the long run has detrimental consequences for the cell.

## Spatial and functional organization of translation

Finally, we investigated the spatial organization of translation in cells (Fig. 4a). It is known that ribosomes translating on the same mRNA can assemble closely in space to form polysomes^12,54,55^. We detected polysomes based on a distance cut-off of 7 nm from the mRNA exit site to the mRNA entry site between neighbouring ribosomes (Extended Data Fig. 11a-e, Supplementary Discussion). Polysomes account for 26.2% of all 70S ribosomes, and two arrangement patterns between neighbouring ribosomes can be defined (Fig. 4b-g), *i*.*e*. the so-called “top-top” (t-t; 78.5%, 4.2 ± 1.4 nm) and “top-back” (t-b; 21.5%, 5.4 ± 1.5 nm) configurations^55^. We also observed various topologies for long polysomes, ranging from loose assembly to tight packing with helix-like configurations (Fig. 4c-f) ^12,55,56^.

**Fig. 4.**
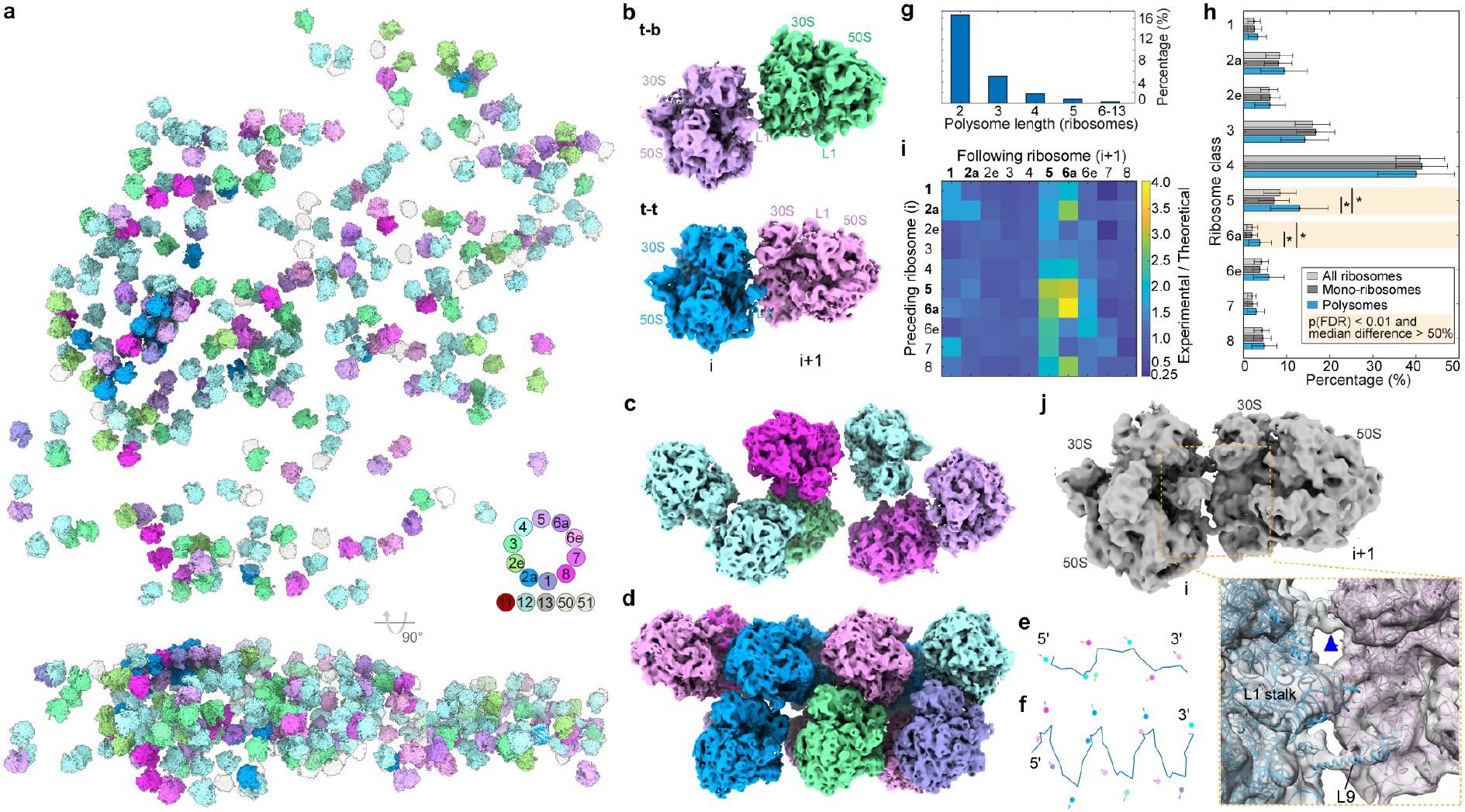
Spatial and functional organization of ribosomes in native cells. **a**, 3D map of ribosomes in a representative untreated cell. Top: x-y view; bottom: orthogonal view. 50S: light grey; 70S colored by translation states as shown in Extended Data Fig. 4, and Fig. 2a (elongation cycle). **b**, Assembly patterns of adjacent ribosomes in polysomes. **c**-**d**, Representative long polysomes of different topologies. **e**-**f**, Putative mRNA paths and nascent chain vectors for **c** and **d** respectively. **g**, Distribution of polysome lengths. **h**, Distribution of elongation states in polysomes compared to all ribosomes and mono-ribosomes. Bars and whiskers are mean and standard deviation across 356 tomograms of untreated cells. Highlighted are states for which the fraction in polysomes differs by more than 50% compared to all ribosomes or mono-ribosomes and FDR-adjusted p-value calculated with Wilcoxon rank sum test < 0.01. **i**, Occurrence frequencies of elongation state in pairs of adjacent ribosomes in polysomes normalized to the theoretical probability of random pairs. States that require elongation factor binding to proceed are 1, 2a, 5 and 6a (bold). **j**, Structure of a di-ribosome within polysomes shows the intervening mRNA density (inset: blue arrowhead) and the stretched L9 of the preceding ribosome.

Whether translation elongation is synchronized or coordinated within polysomes represents a long-standing question^12,54-56^. To address this, we first examined whether the distribution of elongation states differs between the total ribosome population and polysomes. We found that while most states occurred equally frequently in both populations, the fractions of two states prior to EF-G binding (“A*, P/E” and “A/P, P/E”) are more frequent in polysomes (Fig. 4h). We next calculated the frequencies of state pairs between two adjacent ribosomes (preceding *vs*. following) and compared them to theoretical pair frequencies calculated from the bulk distribution (Extended Data Fig. 12a-c). This comparison revealed that the occurrence of states among preceding and following ribosomes is not symmetric: ribosomes in the states requiring elongation factor binding to proceed are more frequent as the following ribosome (Fig. 4i). We statistically validated these differences with a permutation test, and further demonstrated that the asymmetry between preceding and following ribosomes increases as the distance threshold decreases (Extended Data Fig. 12d-h, Methods and Supplementary Discussion). This suggests that local coordination of elongation between adjacent ribosomes in the polysome is likely achieved by obstructing elongation factor binding that is required to proceed to the next state.

To investigate whether local coordination arises from structures specific to ribosomes engaged in polysomes, we performed classification of ribosome pairs in polysomes to better resolve the ribosome-ribosome interface (Fig. 4j, Extended Data Fig. 13a, b). We found that within the interface, the ribosomal protein L9 of the preceding ribosome adopts a stretched conformation and its C-terminal domain protrudes into the elongation factor binding site of the following ribosome (Extended Data Fig. 13c, d). Independent focused classification on L9 of all ribosomes showed that it largely (68.9%) adopts a flat conformation on the ribosome surface, whereas it is stretched in 20.2% of 70S ribosomes (Extended Data Fig. 13e). Ribosomes with a stretched L9 largely overlap with the ribosomes detected as polysomes, especially those with tighter “t-t” arrangement (Extended Data Fig. 13f, g). These results suggest that L9 tends to have a flat conformation in single ribosomes and to adopt the stretched conformation within compacted polysomes. This clarifies previous findings that L9 is stretched in X-ray crystallography structures where the crystal packing recapitulates configurations of compacted polysomes, but is found in the flat conformation in cryo-EM structures of isolated ribosomes^57^. While L9 has been reported to be non-essential, its mutations can cause increased frameshifting and ribosome compaction by one codon^58,59^. We therefore propose that within tightly assembled polysomes, L9 of one ribosome can adopt a stretched conformation that sterically interferes with elongation factor binding to the following ribosome. This local effect can buffer adjacent ribosomes and help to maintain translation fidelity by avoiding direct collision within polysomes.

## Conclusions

Our study demonstrates the potential of cryo-ET in illuminating dynamic processes carried out by macromolecular machines in living cells. The translation elongation cycle retrieved from the single-cell dataset of a genome-reduced bacterium complements current models derived from controlled *in vitro* approaches. The structural and quantitative profiling of the cellular translation machinery presented here improves our understanding of protein biogenesis in cells and elucidates how ribosomes respond to different antibiotic perturbations on the single-cell level. Our investigation of polysomes shows the advantage of in-cell structural biology, which can relate functional states of a molecular machine to its molecular sociology and cellular context. The approaches developed here can be applied to analyse structural dynamics of other cellular processes in the future and will contribute to the construction of cell models at atomic detail.

## Methods

### Cryo-ET sample preparation and data collection

The *Mycoplasma pneumoniae* cell cultivation, sample preparation and data collection were described previously^16^. The three datasets of untreated, Cm-treated and PUM-treated cells were re-processed in Warp/M 1.0.7 (the alpha versions that were officially released as version 1.0.9.)^17,60^. The small dataset acquired with Volta phase plate was processed with the denoising network in Warp 1.0.9 for visualization purpose only (shown in Fig. 1a).

A new dataset of spectinomycin (Spt)-treated cells was collected following the same procedure as before^16^. In brief, spectinomycin (Sigma-Aldrich) at 0.4 mg/ml final concentration was added into the culture medium, 15-20 minutes before plunge freezing. Tilt-series collection with the dose-symmetric scheme^61^ was performed on a Titan Krios equipped with a K3 camera (Gatan), at magnification 81,000x, pixel size 1.053 Å, tilt range -60° to 60° with 3° increment, total dose 120-140 e^−^/Å^2^.

In total, 356 untreated, 65 Cm-treated, 86 PUM-treated, and 70 Spt-treated cellular tomograms were analyzed in this work. Details of cryo-EM data collection, refinement and validation statistics are provided in Extended Data Tables 1-5.

### Image processing, ribosome template matching and map refinement

Pre-processing (motion correction, CTF estimation, dose filtering, tilt-series sorting) was performed in Warp 1.0.9. For the untreated, Cm-treated and PUM-treated datasets, the ribosome coordinates were adopted from previous particle picking^16^. For the Spt-treated dataset, template matching was performed in PyTom^62^, followed by classification to exclude false positives, without manual cleaning. In total, 109,990 untreated, 21,299 Cm-treated, 23,014 PUM-treated, and 13,418 Spt-treated ribosome sub-tomograms were reconstructed in Warp.

3D refinement and classification were performed in RELION 3.0^63,64^. We then used the software M (version 1.0.9) to perform multi-particle refinement of the tilt-series and the averaged structure^17^. Refinement of both geometric (image and volume deformation) and CTF parameters was done in a sequential manner. After M refinement on 70S, we performed focused refinement on 30S and 50S subunits separately to improve the local resolution. Fourier shell correlation (FSC) calculation between independently refined random half subsets, local resolution estimation, and post-processing were done in M.

### Model building for high-resolution ribosome maps

Models for the 30S and 50S subunits were first built based on the maps from focused refinement of Cm-treated ribosomes (EMDB 11998 and 11999). Homology models for ribosomal proteins were generated using SWISS-MODEL^65^ with a *Bacillus subtilis* ribosome as the template (PDB 3J9W), except for L9 (PDB 1DIV and 4V63), L10 (PDB 1ZAV), and S21 (PDB 5MMJ). rRNA sequences (REFSEQ NC_000912) were aligned to the *E. coli* rRNA (PDB 4YBB) using SINA online server^66^. ModeRNA^67^ was used to build homology models for rRNAs based on the previous alignment, with PDB 4YBB as the template. Homology models were rigid-body fitted into the cryo-ET densities using Chimera^68^, followed by iterative refinement using PHENIX real-space refinement^69^ and manual adjustment in Coot^70^. The ribosomal protein L9, L10 and L11 were only fitted as rigid bodies into the map due to the less resolved local density. Focused classification was performed to further resolve structural heterogeneity of L9. The flat conformation of L9 that resides on the ribosome surface was found to be predominant and was built in the model. Sequence extensions for ribosomal proteins S6, L22 and L29 were built *de novo*. Models were validated using MolProbity^71^. FSC curves between the model and the map were also calculated for validation^72^.

### Bioinformatic analysis of ribosomal proteins

Bioinformatic sequence analysis of ribosomal proteins in *M. pneumoniae* and the comparison with *E. coli* homologs, described in details in the Supplementary Methods, were performed as follows: i) protein sequences and RefSeq genome annotation for *M. pneumoniae* strain M129 (ATCC 29342) were downloaded from NCBI; ii) protein sequences were additionally annotated with eggNOG-mapper to obtain COG IDs (Clusters of Orthologous Groups)^73,74^; iii) for the annotated ribosomal proteins, the corresponding COG multiple sequence alignments from representative bacterial species were downloaded from the eggNOG database^75^; iv) as *M. pneumoniae* M129 is not among the representative species, its protein sequences were added to the multiple sequence alignments with MAFFT software^76^; v) for each COG multiple sequence alignment, the number of amino acids in every representative species (including *M. pneumoniae* M129) present at positions before the N-terminus and after the C-terminus positions of *E. coli* K-12 substr. MG1655 were calculated; vi) the presence of N- or C-terminus extensions longer than 20 amino acids were illustrated in iTOL using NCBI taxonomy tree as the basis^77^; vii) for all COGs corresponding to proteins with N- or C-terminus extensions in *M. pneumoniae* that are longer than 20 amino acids, protein disorder and secondary structure were analyzed for all representative strains. The presence of crosslinks or transposon insertions were additionally analyzed.

### Sub-tomogram classification of the translation states of ribosomes

Maximum-likelihood 3D classification was performed in RELION 3.0 with the re-extracted ribosome sub-tomograms after M refinement. A hierarchical and exhaustive classification strategy (Supplementary Discussion) with at least three tiers was used to handle the heterogeneity in the untreated dataset, which is described here in detail and illustrated in Extended Data Figures 4 and 5. A similar procedure was applied for all datasets, and is detailed in Extended Data Figure 8-10.

In the first tier, 109,990 sub-tomograms were classified into 70S and free 50S with a global spherical mask (320 Å diameter). The 24,157 free 50S were further classified into two classes, with a local spherical mask focusing on the ribosome recycling factor binding site. Prior to extensive 70S classification, structural heterogeneity was evaluated by visual inspection, multibody refinement and test classification runs in RELION. Classification setups were extensively tested, including different masks (global 70S mask, local 30S mask, spherical tRNA path mask, solvent tRNA path mask, spherical EF mask, solvent EF mask, *etc*.), initial references (ribosome average, features-less sphere or none), angular search options (global, local or none), and RELION optimization parameters (class numbers 2-16, T values 2-10, iterations 25-40, limit resolution E-step 5-10 Å, *etc*.). To avoid bias, we mostly used a sphere with soft edges as the mask. Local spherical masks covering factors (EFs and/or tRNAs) provided more consistent and stable classification results than bigger overall masks. The class number was made higher than the number of distinct structures that could be retained in one classification job and similar resulting classes were grouped.

In the second tier, the 77,539 identified 70S ribosomes were further classified with a local mask (Extended Data Fig. 4a, mask I) focusing on the tRNA path region, roughly covering the A, P, and E sites. This resulted in four major classes with different tRNA occupancies: “P, E”, “A**, P/E”, “P”, “A, P”. In the following rounds, we could further classify the “A**, P/E” class into “A*, P/E” and “A/P, P/E”. In addition, a 70S class with only one hybrid “P/E” tRNA was classified. Two of the resulting classes were not interpretable: 2,150 particles with dim 30S density that were poorly resolved and 1,484 particles with density near the P site that does not resemble a tRNA.

In the third tier, focused classification with a local mask (Extended Data Fig. 4a, mask II) around the elongation factor and A/T tRNA binding sites was carried out. For the previous class with “P, E” tRNAs, further classification resulted in 1,803 particles without additional density, 3,324 particles with EF-G and 4,634 particles with EF-Tu•tRNA. For the classes with partial and full hybrid tRNAs, sub-classes with EF-G were obtained. For the class with only “P” tRNA, 12,464 particles with additional EF-Tu•tRNA were classified out.

For each classification step, at least three parallel RELION classification jobs with the same or slightly different parameters (local angular search range, class number, T value, *etc*.) were carried out for comparison. The classification job with the most stable result was selected and used for sorting sub-tomograms (Extended Data Fig. 5a-g). After sorting, subsequent validation classification runs were performed for each sorted class to test whether new structures emerge or whether particles were “wrongly” classified. Misclassified particles were relocated to the corresponding class and the validation runs were repeated until convergence. This exhaustive approach was performed for all classification steps. For each of the final classes, refinement and post-processing were done in RELION. Further classification performed with the new refinement results as inputs did not generate any new classes.

### Model building and comparison of the ribosome translation states

To obtain models for ribosome classes, the 30S and 50S models built above were used as starting models for flexible fitting. Homology models of EF-G and EF-Tu were made by SWISS-MODEL with PDB entry 4V7D and 4V5L as the template, respectively. For tRNA homology models, the tRNA in an *E. coli* ribosome structure (PDB 4V7C) was used as the template and mutated to the sequence of *M. pneumoniae* Phe-tRNA (REFSEQ NC_000912). The mRNA and nascent peptide were built using PDB 3J9W as the template. The homology model of the ribosome recycling factor was built using PDB 1EH1 as the template. For each class, the starting models were first rigid-body fitted into the density using Chimera and then flexible fitting was done using Namdinator^78^. Validation was performed as described above.

For measuring 30S body rotations, structures of all classes were first aligned to the 50S subunit and then the rotation angles were estimated with a pivot point near the nucleotide 11 of the 16S rRNA. With the perspective from the solvent side of the 30S subunit, positive numbers equal counter-clockwise rotations and negative numbers equal clockwise rotations. For measuring 30S head rotations, all class structures were first aligned based on the 30S body and then the rotation angles were determined with the axis near the 30S neck. For both rotations, the angles in the “A, P” class were defined as 0°. To describe the L1 dynamics, the distance between the mass center of the L1 stalk (near nucleotide 2,181 of the 23S rRNA) and a fixed point near the center of the classical E site (determined based on the “P, E” class) was measured after aligning all classes based on the 50S subunit (excluding the L1 stalk).

### Spatial analysis of ribosomes and polysomes

Spatial mapping of ribosomes within one cellular tomogram was achieved by projecting back the ribosome structures into the tomograms, with coordinates determined by template matching and shifts and rotations determined by RELION refinement. The projection was performed using the TOM toolbox^79^ after Euler angle format conversion, at 4x or lower binning (voxel size 6.8Å). To calculate the ribosome concentration, we first estimated the cellular volume covered in the tomogram and then divided the total number of detected ribosomes by the volume.

Detection of the polysomes was based on both position and orientation information (Extended Data Fig. 11), using a homemade script in MATLAB 2016b. The positions of mRNA entry and exit sites for all 70S ribosomes were calculated based on the rotations and shifts determined during RELION refinement. The distances from the mRNA exit site of one ribosome (as the preceding one) to the mRNA entry sites of all neighbouring ribosomes (as the potential following ribosome) were calculated. A distance threshold of 7 nm was used to define whether two ribosomes belong to the same polysome. The calculation and polysome definition were done for all 70S ribosomes. A unique identifier was assigned to each polysome, as well as the sequential number for all ribosomes within the polysome.

The distribution of relative positions of adjacent ribosomes within the polysome, *i*.*e*. the position of the following ribosome (i+1) relative to the preceding ribosome (i), was analyzed after normalizing the relative position vector with rotations for the preceding ribosome determined during refinement. The distribution of relative rotations of two adjacent ribosomes was represented as three Euler angles (ψ, θ, φ in XYZ system) of the following ribosome. Using the k-means clustering function in MATLAB, we determined two major arrangement configurations for adjacent ribosomes within the polysome, *i*.*e*. how the following ribosome rotates relative to the preceding ribosome. These two configurations are identical to the previously reported “top-top” and “top-back” configurations^55^.

To further refine the ribosome-ribosome interface in polysomes, classification was performed with sub-tomograms extracted with a large box size that can accommodate two ribosomes. After refinement on the preceding ribosome (i), classification without refinement was performed with a local mask focusing the following ribosome (i+1). Models for the resulting classes were built with rigid-body fitting of previously obtained ribosome models and the L9 homology model with the stretched conformation (PDB 4V63).

### Statistical analysis of translation elongation states in polysomes

To compare the experimental elongation state frequencies in polysomes with theoretical frequencies, the distributions of frequencies of each elongation state calculated across 356 tomograms of the untreated cells were compared between polysomes and all ribosomes, and between polysomes and mono-ribosomes by calculating the fold change between distribution medians. Significance was assessed with a two-sided Wilcoxon-Mann-Whitney test using the ranksum function in MATLAB 2019b. The theoretical frequency of each ribosome pair was calculated as the product of the overall frequencies of the ribosome classes for the preceding ribosome (i) and the following ribosome (i+1). Experimental polysome pair frequencies were calculated by summarizing the numbers of all ribosome pairs engaged in polysomes, and dividing these numbers by the total number of pairs. Experimental and theoretical pair frequencies were compared by calculating the fold change per pair.

Permutation analysis was performed to test the significance of different occurrence frequencies of the experimental pairs compared to the theoretical. The ribosome pair sequences in polysomes were represented as a matrix with single polysomes as rows and ribosome positions as columns, where each matrix cell contains the ribosome class representing its elongation state. Columns with positions beyond each ribosome’s length were assigned to NaN (not-a-number). All polysomes across all tomograms were combined in one matrix (8,641 polysomes in total). For permutation analysis, all elements of the polysome matrix were randomly shuffled for 10,000 times with the randperm function in MATLAB. For each row, the elements were sorted so that the ribosome classes are in the front columns followed by NaNs occurring in the same row after shuffling. Rows with less than two ribosomes in the sequence were deleted. Shuffled ribosome pair frequencies were calculated the same as experimental pair frequencies. Permutation *p*-value for each ribosome pair was calculated as the minimum between the number of permutations in which the pair frequency was less or equal to the experimentally observed frequency divided by the total number of permutations, and the one minus this value. Permutation *p*-values were adjusted for multiple hypotheses testing with the Benjamini-Hochberg procedure using the mafdr function in MATLAB with parameters (‘bhfdr’, ‘true’).

For polysome distance threshold analysis, matrices were created from the full ribosome dataset by varying the distance threshold to the nearest neighbour in the range from 3 to 10 nm. The distances for polysome definition were calculated between the mRNA exit site of one ribosome and the mRNA entry site of another ribosome as described above. The fractions of elongation states in ribosome pairs were calculated as for the original ribosome set (with the threshold of 7 nm). For each distance threshold and each ribosome class, the ratio between the number of pairs in which the preceding ribosome has this class and the number of pairs in which the following ribosome has this class was calculated.

### Single cell clustering analysis

The distribution of 70S ribosome classes identified in the translation elongation phase for all four datasets (356 untreated cells, 65 Cm-treated cells, 70 Spt-treated cells, 86 PUM-treated cells) were used for clustering analysis. As each tomogram covers most of one cell, the sub-tomograms in each class could be further separated by which cell/tomogram they belong to. Classes in the antibiotic-treated cells were assigned with the class identifiers according to the closest classes in the untreated dataset. The percentages of different classes within each cell/tomogram were calculated and these numbers were used as inputs for clustering analysis. Hierarchical clustering analysis was done using the clustergram function in MATLAB 2016b.

Structure visualization, preparations for figures and movies were done in Chimera^68^ and ChimeraX^80^.

## Data availability

Maps were deposited in the Electron Microscopy Data Bank (EMDB) under accession numbers: 13234, 13272, 13273, 13274, 13275, 13276, 13277, 13278, 13279, 13280, 13281, 13282, 13283, 13284, 13285, 13286, 13410, 13411, 13412, 13413, 13414, 13431, 13432, 13433, 13434, 13435, 13436, 13445, 13446, 13447, 13448, 13449, 13450, 13451, 13452, 13287, 13288, 13289. Models were deposited in the Protein Data Bank (PDB) under accession numbers: 7OOC, 7OOD, 7P6Z, 7PAH, 7PAI, 7PAJ, 7PAK, 7PAL, 7PAM, 7PAN, 7PAO, 7PAQ, 7PAR, 7PAS, 7PAT, 7PAU, 7PH9, 7PHA, 7PHB, 7PHC, 7PI8, 7PI9, 7PIA, 7PIB, 7PIC, 7PIO, 7PIP, 7PIQ, 7PIR, 7PIS, 7PIT. Detailed information for all structures is available in the Extended Tables. Maps and atomic models used from previous studies were obtained from EMDB (11998, 11999) and PDB (3J9W, 1DIV, 4V63, 1ZAV, 4YBB).

## Code availability

The code and associated data for bioinformatics analysis of ribosomal protein extensions, and statistical analysis of polysome sequences were deposited under GitHub repository https://github.com/mszimmermann/mycoplasma_ribosome.

## Acknowledgements

We thank Ievgeniia Zagoriy, Lukas Adam, Felix Weis and Wim Hagen for technical support, and Luis Serrano for providing the *M. pneumoniae* transposon library and proteomics data. We are grateful to Christian Spahn, Arjen Jakobi and Stefan Pfeffer for fruitful discussions and critical reading of the manuscript. J.R. acknowledges the Deutsche Forschungsgemeinschaft (project no. 426290502) and the Wellcome Trust Senior Research Fellowship (103139). J.M. acknowledges funding from the EMBL and the European Research Council starting grant (3DCellPhase^-^760067).

## Author contributions

L.X. and J.M. designed the study; L.X. collected and processed the cryo-ET data, and performed structural analysis; S.L. and L.X. built the atomic models; M.Z.K. performed the bioinformatic and statistical analysis; D.T. and P. C. provided software support and advice on data interpretation; L.X. and J.M. wrote the manuscript with inputs from all authors.

## Competing interests

The authors declare no competing interests.

## Additional information

Supplementary Information is available for this paper.

## Supplementary Information

### Supplementary Methods

The detailed procedure for the bioinformatic analysis of ribosomal proteins for identification and characterization of sequence extensions was provided as below. RefSeq genome annotation and protein sequences were downloaded from NCBI for the *M. pneumoniae* M129 GCF_000027345.1 assembly from https://ftp.ncbi.nlm.nih.gov/genomes/all/GCF/000/027/345/GCF_000027345.1_ASM2734v1/ (accessed 21/04/2021). File GCF_000027345.1_ASM2734v1_genomic.gff was used to identify ribosomal proteins annotated by RefSeq. File GCF_000027345.1_ASM2734v1_protein.faa with protein sequences in FASTA format was used as input to the online tool eggNOG-mapper v2 which uses precomputed eggNOG v5.0 clusters and phylogenies for fast orthology assignment^73,75^. The annotation tables by RefSeq and eggNOG-mapper were merged by protein ID. In total, 52 proteins were annotated as ribosomal and mapped to 51 unique COGs (two proteins, WP_010874426.1 and WP_010874827.1, mapped to the same COG0267 corresponding to ribosomal protein L33).

Multiple sequence alignments for proteins from representative bacterial species were downloaded from eggNOG v5.0 database using eggNOG API. For each of the 51 COG IDs mapping to ribosomal proteins in *M. pneumoniae*, the trimmed alignments were loaded from http://eggnogapi5.embl.de/nog_data/text/trimmed_alg/COG_ID. To add ribosomal proteins from *M. pneumoniae* M129 to these multiple alignments, the sequence of each of the proteins was saved in a separate fasta file, and the MAFFT software v7.475 was used in the add mode for each alignment as follows: mafft --reorder --add protein.fasta --auto trimmed_alg_COG.fa > output_file^76^. For COG0627, the protein WP_010874827.1 from *M. pneumoniae* was used for alignment.

For each of the ribosomal COGs, the amino acid positions of the protein from *E. coli* strain K-12 substr. MG1655 was taken as a reference, and for each protein from other species the number of amino acids before the N-terminus and after the C-terminus locations in *E. coli* was calculated. Proteins that had > 20 amino acids N- or C-terminus extensions in *M. pneumoniae* M129 were considered for further analysis (2 proteins with N-terminus extensions and 9 proteins with C-terminus extensions). Ribosomal protein S3 (COG0092), which has a 17 amino acid extension at the C-terminus, was also retained for further analysis.

NCBI bacterial taxonomy file new_taxdump.zip was downloaded from https://ftp.ncbi.nlm.nih.gov/pub/taxonomy/new_taxdump/ (version on 24/04/2021). The file names.dmp was used to map NCBI IDs to taxonomy names. Tree was reconstructed from the file taxidlineage.dmp in Python v3.7.7 with ETE3 Toolkit v3.1.2^81^. EggNOG protein IDs containing NCBI species IDs were directly mapped to the NCBI tree nodes. The presence of ribosomal protein extensions was saved as tables and converted to iTOL format with table2itol utility (https://github.com/mgoeker/table2itol) in R v3.6.1. The tree was visualized with iTOL v6^77^.

Sequences for 11 ribosomal proteins in *M. pneumoniae* and their orthologs in other species were further analyzed with disorder and secondary structure prediction tools. For disorder prediction, all protein sequences from the multiple sequence alignment files were saved in FASTA format without gaps. IUPred2A tool in Python v3.7.7 was run with the input parameter “long” for each protein sequence^82^. The number of disordered amino acids and disorder length in extended regions were calculated based on the position of the extended region relative to *E. coli*, and IUPred2A score of 0.5 was used as the disorder threshold.

All protein sequences from the multiple sequence alignment files were saved as separate fasta files without gaps in the sequence. JPred prediction tool^83^ was used to predict secondary structure of each protein using the jpredapi utility and JPred-big-batch-submission utility for a large number of submissions (https://github.com/fabianegli/JPred-big-batch-submission). The number of helices in extended regions were calculated based on the position of the extended region relative to *E. coli* protein.

For 11 ribosomal proteins in *M. pneumoniae* with N- or C-terminus extensions compared to *E. coli*, the positions of crosslinks linking to the same protein or a different protein were mapped according to the amino acid crosslink positions reported previously^16^.

To analyze transposon insertions in *M. pneumoniae* ribosomal proteins with extensions, the supplementary materials from a previous publication^19^ was used. All the files in the “SupplementaryData1_fastqins_processed.zip folder” (files with extension ‘.qins’ but not ‘_fw.qins’ or ‘_rv.qins’) were concatenated to obtain a list of nucleotide coordinates of transposon insertions in the *M. pneumoniae* M129 NC_000912.1. The transposon insertion locations in the 11 ribosomal proteins were selected based on their genomic locations as per RefSeq annotations (GCF_000027345.1_ASM2734v1_genomic.gff). Nucleotide locations were converted to amino acid locations by calculating the difference between each transposon location and the start coordinate of the corresponding gene and dividing the number of nucleotides by three.

**Supplementary Table 1.** Statistics of the eleven identified ribosomal protein with extensions across bacteria.

| Protein | S2 | S3 | S5 | S6 | S18 | L3 | L4 | L22 | L23 | L29 | L31 |
| --- | --- | --- | --- | --- | --- | --- | --- | --- | --- | --- | --- |
| Length of extension in *M. p.* (amino acids) | 46 | 17 | 57 | 74 | 30 | 74 | 50 | 49 | 139 | 47 | 25 |
| Number of disordered residues (score>0.5) in *M. p.* | 29<br>(63%) | 17<br>100% | 57<br>100% | 67<br>(90%) | 23<br>(76%) | 68<br>(91%) | 2<br>(4%) | 46<br>(93%) | 110<br>(79%) | 11<br>(23%) | 8<br>(32%) |
| Number of predicted helices in *M. p.* | 19<br>(41%) | 0<br>(0%) | 1<br>(1%) | 23<br>(31%) | 0<br>(0%) | 5<br>(6%) | 15<br>(30%) | 23<br>(46%) | 0<br>(0%) | 28<br>(59%) | 0<br>(0%) |
| Number of predicted beta sheets in *M. p.* | 0<br>(0%) | 0<br>(0%) | 0<br>(0%) | 0<br>(0%) | 2<br>(6%) | 2<br>(2%) | 10<br>(20%) | 0<br>(0%) | 0<br>(0%) | 0<br>(0%) | 0<br>(0%) |
| Number of species with extensions (out of 4,396) | 1,956<br>(44%) | 851<br>(19%) | 895<br>(20%) | 50<br>(1%) | 396<br>(9%) | 579<br>(13%) | 3,438<br>(78%) | 212<br>(4%) | 44<br>(1%) | 159<br>(3%) | 138<br>(3%) |
| Number of species with >50% disordered residues in extensions | 1,618<br>(82%) | 840<br>(98%) | 890<br>(99%) | 16<br>(32%) | 200<br>(50%) | 501<br>(86%) | 55<br>(1%) | 193<br>(91%) | 20<br>(45%) | 150<br>(94%) | 113<br>(81%) |
| Number of species with >50% helices in extensions | 50<br>(2%) | 0<br>(0%) | 0<br>(0%) | 2<br>(4%) | 0<br>(0%) | 0<br>(0%) | 0<br>(0%) | 84<br>(39%) | 26<br>(59%) | 42<br>(26%) | 17<br>(12%) |
| Number of species with >50% beta sheets in extensions | 0<br>(0%) | 0<br>(0%) | 0<br>(0%) | 0<br>(0%) | 0<br>(0%) | 0<br>(0%) | 0<br>(0%) | 0<br>(0%) | 0<br>(0%) | 0<br>(0%) | 0<br>(0%) |

*M. p.*: *Mycoplasma pneumoniae*

## Supplementary Discussion

### Atomic ribosome model reveals divergent protein extensions

Refinement of the 18,987 sub-tomograms containing 70S ribosomes from 65 Cm-treated cells, and refinement of the 77,539 sub-tomograms from 356 untreated cells both resulted in 70S ribosome maps with a nominal resolution of 3.5 Å (Extended Data Fig. 1a-f). Local resolutions of the 50S subunit in both datasets are at the Nyquist limit of 3.4 Å (Extended Data Fig. 1b-j). For the 30S subunit, focused refinement of the Cm-treated ribosomes generated higher resolution (3.7 Å) compared to the untreated (Extended Data Fig. 1b-j), because of the reduced structural flexibility upon Cm binding. The better resolved densities from the Cm-treated dataset were used to build atomic models for the 30S and 50S subunits separately (Extended Data Fig. 1g-k). These were then used as initial models to build models for the 70S ribosome.

Extensions of the eleven ribosomal proteins are all solvent-exposed on the ribosome surface and positioned away from the active sites (Fig. 1c, Extended Data Fig. 2e). They are therefore not expected to be directly involved in the translation process. However, the ribosomal proteins bearing these extensions are located near important interaction interfaces of the ribosome. Our previous in-cell crosslinking mass spectrometry results^16^ also indicate that the extensions are crosslinked to different proteins (Extended Data Fig. 3a). Ribosomal proteins S2, S3 and S5 are located near the mRNA entry site on the ribosome, and have been suggested to form the main contact interface with the RNA polymerase during transcription-translation coupling in *M. pneumoniae*^16^. S6 resides near the interface between the two ribosomal subunits, but its long C-terminal extension can extend towards the mRNA entry site. S6 is crosslinked to the delta subunit of the RNA polymerase (Extended Data Fig. 2f-g). These results suggest that the extensions of 30S proteins may have functional roles near the mRNA entry site of the ribosome. On the 50S subunit, three (L22, L23 and L29) of the four ribosomal proteins surrounding the nascent peptide tunnel have extended sequences (Fig. 1c, Extended Data Fig. 2e, 3a). The long helices formed by the extensions of L22 and L29 mainly interact with peripheral rRNA. All three extensions are over 30 Å away from the nascent peptide exit site (Fig. 1c, Extended Data Fig. 2e), and do not appear to be able to interact with the nascent peptide during translation. Although functions of these extensions remain largely elusive, a transposon mutation screen^19^ in *M. pneumoniae* shows that their disruption affects cellular fitness or survival (Extended Data Fig. 3a).

Our bioinformatic analysis revealed that 1% to 78% out of the 4,396 tested representative bacteria strains have ribosomal protein extensions in comparison to *E. coli* K-12 (Extended Data Fig. 3, Supplementary Table 1). The largest frequency of extensions (78%) was detected in L4, including the extension in *M. pneumoniae* M129, although the sequence lengths are comparable between species. This possibly reflects the fact that the sequence at the C-terminus of L4 is different in Gammaproteobacteria compared to other bacteria, which results in multiple sequence alignment where many bacteria have a C-terminus extension compared to *E. coli*. For the other ten ribosomal proteins with extensions, the extensions are at least 20 amino acids longer in *M. pneumoniae*, compared to both *E. coli* and *B. subtilis*. These protein extensions, however, are not specific to any phylum or sub-groups of bacteria (Extended Data Fig. 3b). Within the Tenericutes phylum or the Mollicutes class to which *M. pneumoniae* belongs, the extensions appear to be more frequent than other bacteria. Moreover, extensions in other bacteria, such as *Mycobacterium smegmatis*^84-86^, are not limited to the eleven ribosomal proteins found in *M. pneumoniae*. The observation that *M. pneumoniae* has many ribosomal proteins with extensions being a genome-reduced bacterium that lives a strict parasitic lifestyle as a human pathogen^87^, is reminiscent of reports that parasitic protozoans also exhibit extensive ribosomal protein extensions compared to other eukaryotes^88^. Structure determination of ribosomes from non-model species will be necessary to understand evolution and diversity of ribosomes in adaptation to different environments.

### Ribosome classification for probing the translation process in cells

The structural dynamics of ribosomes along their functional trajectories, as well as the distribution of these states, are expected to be preserved in a close-to-native condition in frozen-hydrated cells. The vitrification time^89^ (0.1 - 0.2 microsecond) is much faster than any known translation steps^8,35,36^ (at least at millisecond scale). *M. pneumoniae* cells in the fast-growing phase were directly grown on carbon-coated gold grids at 37 °C in the rich medium, minimising any potential perturbation prior to plunge freezing. Vitrification was done with a manual plunger without a temperature controller. The cells on the grid were exposed to room temperature (20 - 25 °C) during the blotting step (∼3 seconds). We cannot exclude the possibility that this short exposure led to a slight temperature change to cells on the grid.

A hierarchical and exhaustive classification procedure was designed to mitigate potential variations associated with individual RELION classification jobs (Methods, Extended Data Fig. 4, 5), and to ensure the proportions of classified translation states are representative of their distribution inside cells. We further validated unbiased representation of 70S ribosome localization and the robustness of the classification, by plotting the class distribution against template matching cross-correlation coefficients (Extended Data Fig. 5h, i). The extracted sub-tomograms retrieved the vast majority (>90%) of 70S ribosomes and a subpopulation (∼50%) of free 50S in the cellular tomograms (Extended Data Fig. 5h). The analysis shown that there is no discrimination for different 70S classes due to template matching (Extended Data Fig. 5i). Thus, the obtained structures and their distribution are the good approximation of the 70S ribosome population in native *M. pneumoniae* cells. We recognize that rare states, such as those of the initiation and termination phases, may not be identified as distinct classes^6^. The classes that were determined represent states that are more populated and structurally distinct. They are possibly averages of ensembles of several close states^35^.

The sub-tomograms in the most populated “A, P” class could not be further classified to distinguish states before and after peptidyl transfer. Although the nascent peptide density around the A site appears to be stronger compared to the P site, it is impossible to determine the exact state from the density alone (Fig. 2b). No class with all three A-, P- and E-site tRNAs was resolved, different from a number of *in vitro* structures^90-92^. Although our result does not rule out the possibility that ribosomes with all three A-, P- and E-site tRNAs exist in cells, they may represent transient intermediates with low occurrences. This assumption agrees with a previous single-molecule study showing that ribosomes are rarely occupied (1.7%) by three tRNAs during active translation^31^. We could not identify any class with EF-G binding to a non-rotated 30S, consistent with a previous publication showing that this represents a less favourable state^30^. For the two classes with EF-Tu, the blurred local density indicates the motion of EF-Tu during decoding and accommodation^92,93^. Further classification with different focused masks and setups, however, did not result in any distinct sub-classes.

In addition to the ten elongation classes, we captured five classes that may represent states in other translation phases (Extended Data Fig. 4a). The “P/E” class is a rotated 70S ribosome with only one hybrid P/E-site tRNA (Extended Data Fig. 7k), and is expected to be an intermediate state during ribosome recycling^6,94-96^. No other 70S class in the recycling phase was detected. The “P/E” state thus may indicate one of the rate-limiting steps during ribosome recycling, *i*.*e*. the next step of EF-G or ribosome recycling factor (RRF) binding to the ribosome in “P/E” state^21^. The “Dim30S-50S’’ class has a slim 30S subunit and shows no clear tRNA density (Extended Data Fig. 4a, class ⑬). A class of 70S ribosomes was found to have unclear density near the P site (Extended Data Fig. 4a, class ⑫). The densities for these two classes were of too low resolution to be unambiguously interpreted. The free 50S could be further classified into two classes: with and without RRF (Extended Data Fig. 4a). For the “50S-RRF” class, the density for the three helices of RRF’s domain I is clearly resolved, but domain II is blurred (Extended Data Fig. 7l).

### Translation landscapes in *M. pneumoniae*

We consider the frozen-hydrated cells to represent a snapshot of the steady state of the non-equilibrium open system of a living cell^97^. For translation elongation, the steady state is represented by the relatively stable occurrence frequencies of the elongation intermediates across 356 untreated *M. pneumoniae* cells (Fig. 3a, e). *M. pneumoniae* cells are known to have slow growth rate and long duplication time (as long as 8 hours)^11,87,98,99^, potentially contributing to the observation of relatively low variations between different cells (Fig. 4e). The reported numbers of ribosomes per *M. pneumoniae* cell vary from 80 to 863^11,100,101^. Our work provides an estimation of 300 to 500 ribosomes per cell. Through dividing these number by the cellular volume estimated from tomograms, we calculated the averaged concentration of 70S ribosomes in *M. pneumoniae* to be 7,400 ± 1,600 µm^-3^. This is considerably lower than the numbers reported in other bacteria, *e*.*g*. 15,000 µm^-3^ in *Spiroplasma melliferum*^102^, and 27,000 - 60,000 µm^-3^ in *E. coli*^103^. Additionally, concentrations of translational factors in *M. pneumoniae* (EF-G, ∼10,000 µm^-3^; EF-Tu, ∼40,000 µm^-3^; total tRNAs, ∼14,500 µm^-3^)^104,105^ are lower compared to *E. coli* (EF-G, ∼75,000 µm^-3^; EF-Tu, ∼500,000 µm^-3^; total tRNAs, ∼500,000 µm^-3^)^106-108^. The different intracellular concentrations of ribosomes and translation factors, which constitute the translation machinery, may contribute to different translation landscapes across bacteria, and other organisms.

### Spatial analysis of ribosomes and polysomes

A tomogram in our data can cover 70% to 90% of the volume of a *M. pneumoniae* cell. The functional state determined for each ribosome can be spatially mapped into the cellular volume (Fig. 4a, b, Extended Data Fig. 11a). To determine the most appropriate criterion for polysome detection, we first calculated the distance distribution for all neighbouring ribosomes and found a peak corresponding to polysomes (Extended Data Fig. 11d). Next, we tested different distance thresholds, ranging from 2 to 10 nm, to define polysomes (Extended Data Fig. 11e). The distance criterion of 7 nm provided the best recovery rate and detection accuracy. The percentage of ribosomes detected as polysomes (26.2%) is consistent with the literature that polysomes are often found at relatively lower abundance (∼30%) in bacteria^109,110^. Most polysomes detected in *M. pneumoniae* cells consist of two to three ribosomes, in agreement with a previous work that monitored real-time translation in living cells using fluorescent microscopy^111^. The average distance from the mRNA exit site of the preceding ribosome (i) to the mRNA entry site of the following ribosome (i+1) is 4.2 ± 1.4 nm for “t-t” pairs (9,100 ribosome pairs), and 5.4 ± 1.5 nm for “t-b” pairs (2,491 ribosome pairs). The “t-t” configuration poses tighter packing of ribosomes compared to the “t-b” configuration.

### Coordination of translation elongation within the polysome

To probe possible coordination of translation elongation in polysomes, we analysed the frequencies of different state combinations in sequential ribosome pairs in polysomes. The experimental ribosome state pair frequencies show similar global distribution to either the theoretical state pair frequencies calculated from individual state frequencies, or the state pair frequencies of shuffled polysomes (Extended Data Fig. 12a-c). This suggests that on the overall level there is no synchronization of elongation states within polysomes (also see Fig. 4c, d). The shuffled pairs were calculated from the shuffled polysome matrices (Methods, Extended Data Fig. 12d), and represent the distribution of pair frequencies as expected if polysomes assemble randomly. The polysome matrix shuffling procedure allowed us to estimate statistical significance of the differences between the experimental and the shuffled state pair frequencies with a permutation test, where the *p*-value represents the fraction of random permutations in which the state pair frequency was smaller (or larger) than the experimentally observed one (Extended Data Fig. 12d, e). The results demonstrated that there is a number of pairs for which the experimental value is significantly different from the shuffled value, though most of these differences are between low-frequency pairs. Notably, the majority of these pairs (20 out of 22) include ribosome states that require elongation factor binding to proceed in the elongation cycle (states 1, 2a, 5, 6a). For these four states, the number of pairs in which the ribosome in this state is a preceding ribosome is significantly lower than expected, and the number of pairs in which the ribosome in this state is a following ribosome is significantly higher than expected (Extended Data Fig. 12e, f). Moreover, we found that this asymmetry in state distribution for two adjacent ribosomes in the polysome becomes more pronounced as the distance between the two ribosomes decrease (Extended Data Fig. 12g). Indeed, for polysomes defined with lower distance thresholds, the ratio of the number of pairs in which the state is a preceding ribosome to the number of pairs in which the state is a following ribosome decreases for states 1, 2a, 5 and 6a, but increases for the other states (Extended Data Fig. 12h). These observations suggested that elongation within polysomes represents a local coordination mechanism that requires close proximity between two neighbouring ribosomes.

Ribosomal protein L9 is conserved across bacteria, but is absent in archaea and eukaryotes^58,112^. L9 has been demonstrated to be flexible, with two RNA binding domains connected by a long helix^57,113,114^. In 70S ribosome crystals, L9 competes with translational GTPases on the following ribosome, preventing crystallization of 70S ribosomes together with these elongation factors^115^. In our structures with better resolved ribosome-ribosome interface, the C-terminal domain of L9 is in contact with the 16S rRNA of the following ribosome (Fig. 4j, Extended Data Fig. 13a, b), similar to the arrangement in crystals. The stretched L9 can generate a steric clash with both EF-G and EF-Tu binding to the following ribosome (Extended Data Fig. 13c, d). The overrepresentation of states prior to EF-G binding is more significant than that of states prior to EF-Tu binding. EF-G mediated translocation is the step where the following ribosome (i+1) moves one codon forward along mRNA and is thereby more likely to directly interact with the stretched L9 of the preceding ribosome (i). In all resolved polysome structures, the following ribosome does not appear to block the L1 stalk opening or tRNA disassociation in the preceding ribosome (Fig. 4j, Extended Data Fig. 13a, b).

### Effects of antibiotics on the translation machinery

Although the exact concentrations of antibiotics inside cells cannot be determined, the drugs were applied at concentrations much higher than the reported saturating concentrations (Cm, Kd = 2 - 6 µM; Spt, estimated saturating concentration = 0.1 mM; PUM, IC50 = 0.1 µM)^45,49,116^. In Cm-treated cells, the Cm density in the three minor classes could not be resolved, possibly due to low resolutions of the maps (Extended Data Fig. 8d). The percentage of ribosomes detected as polysomes is 20.63%, similar to the number in untreated cells. This is in agreement with the observation that Cm treatment has limited effect on polysomes^110^. The average distance from the mRNA exit site to the mRNA entry site for detected polysomes is 3.2 ± 1.5 nm for “t-t” pairs (1,959 pairs), and 5.3 ± 1.1 nm for “t-b” pairs (186 pairs). In Spt-treated cells, the map shows the Spt molecule is in close proximity to the amino acid residue K81 (equals to K26 in *E*.*coli*) on loop 2 of the ribosomal protein S5 (Fig. 3d). This agrees well with reports showing that mutations in the loop 2 regions confer resistance to Spt and affect translation fidelity^117,118^. Whether Spt is bound to ribosomes of the other four classes could not be unambiguously determined due to low resolutions. The percentage of ribosomes detected as polysomes is 12.83%. The average distance from the mRNA exit site to the mRNA entry site for detected polysomes is 4.3 ± 1.5 nm for “t-t” pairs (580 pairs), and 5.1 ± 1.2 nm for “t-b” pairs (326 pairs).

PUM is a nucleoside analogue that specifically competes with UTP for the NTP addition site in RNA polymerases^49^. We have previously shown that the stalled RNA polymerase acts as a physical barrier for the leading ribosome in the transcription-translation coupling complex (stalled expressome)^16^. In contrast to the ribosome-specific antibiotics Cm and Spt, the percentage of detected polysomes under PUM treatment was considerably lower (7.73%). The average distance from the mRNA exit site to the mRNA entry site for detected polysomes was found to be exceptionally small for “t-t” pairs (2.3 ± 0.9 nm; 635 ribosome pairs), and 4.9 ± 1.4 nm for “t-b” pairs (16 ribosome pairs). Taken together, these results suggest that both the elongation state distribution and the spatial organization of the translation machinery are completely reshaped upon different antibiotic treatment.

## Notes

### Competing Interest Statement

The authors have declared no competing interest.

