## Supplementary Figures 1-13 for "Visualizing translation dynamics at atomic detail inside a bacterial cell"

### Extended Data

#### Extended Data Figures

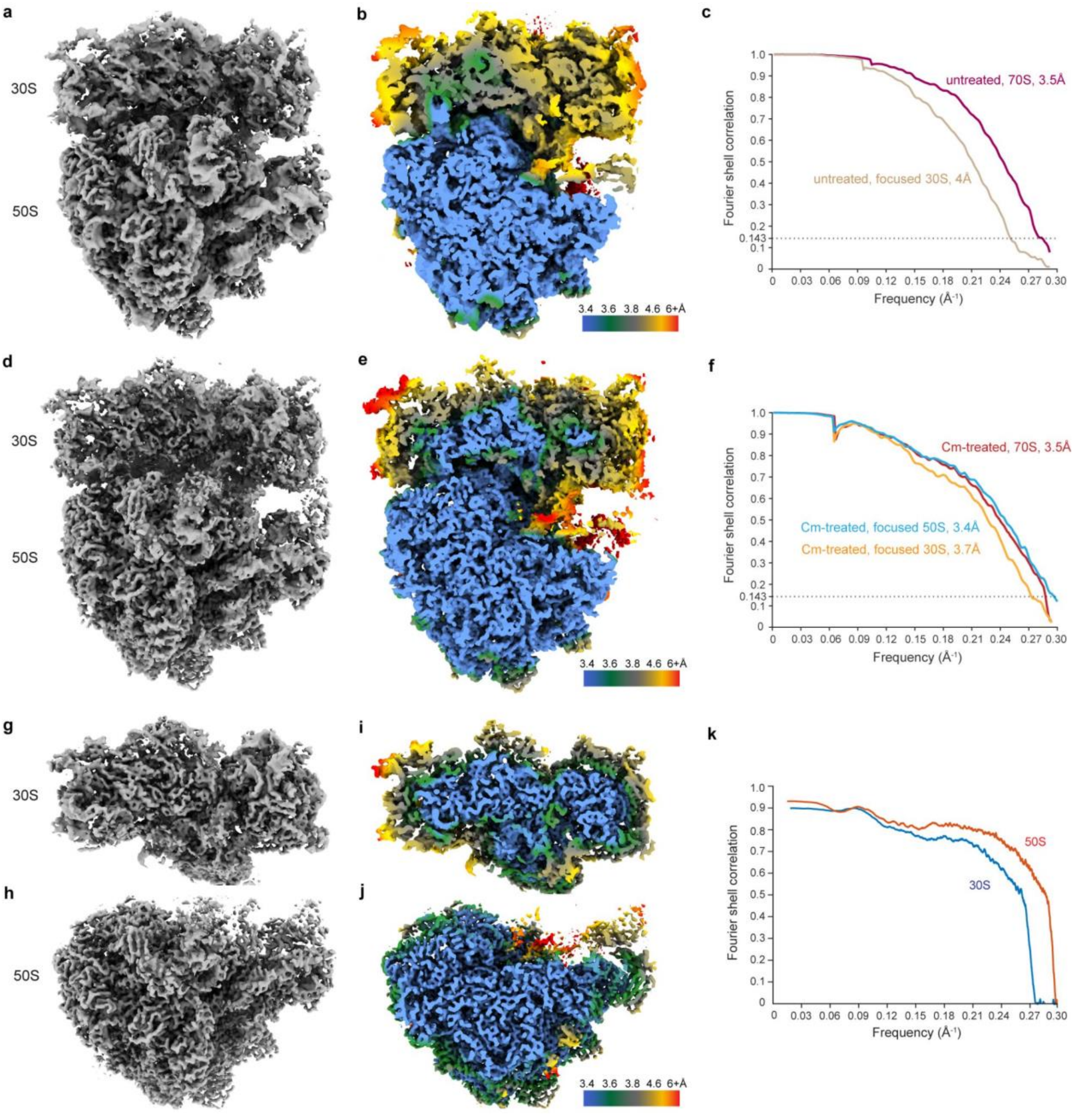


**Extended Data Fig. 1 | *M. pneumoniae* in-cell ribosome maps.**

**a**, 70S ribosome map determined from 77,539 sub-tomograms from 356 untreated *M. pneumoniae* cells. **b**, Map colored by local resolution. **c**, Fourier shell correlation (FSC) curves for global 70S and focused 30S refinement, and the reported resolution value at 0.143 FSC. The Nyquist limit for the data is 3.4 Å. **d**, 70S ribosome map determined from 18,987 sub-tomograms from 65 Cm-treated cells. **e**, Map colored by local resolution. **f**, FSC curves of the Cm-treated 70S ribosome, and of focused refinements on 30S and 50S respectively. **g-h**, Focused refined 30S and 50S maps from the Cm-treated dataset. **i-j**, The corresponding local resolution maps. Atomic models for 30S and 50S were first built based on maps in **g** and **h**. **k**, Model-to-map FSC curves for 30S and 50S respectively.

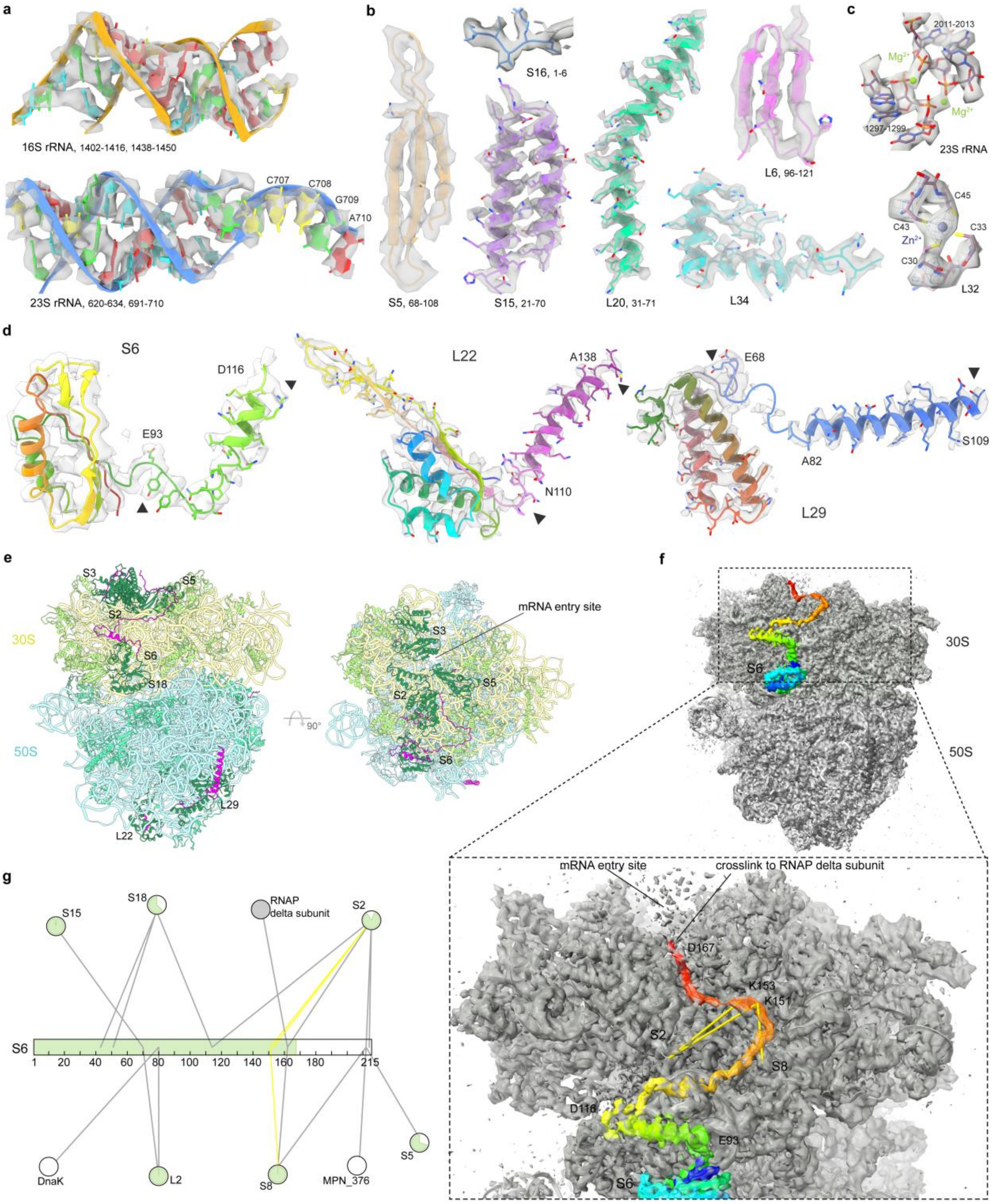


**Extended Data Fig. 2 | Structural features of the *M. pneumoniae* ribosome.**

**a-b**, Examples of regions of the atomic model fitted into the density map for ribosomal RNAs and proteins. **c**, Density corresponding to ions are also observed. **d**, Sequence extensions of ribosomal proteins S6, L22 and L29 form secondary structures (between arrowheads) that were clearly resolved in the map. **e**, The model showing ribosomal proteins with sequence extensions (dark green). The extensions built in the model are highlighted in magenta. In addition to the helix-forming sequences of S6, L22 and L29, some loops can be traced, especially the long C-terminal loop of S6. **f**, Whole-cell crosslinking mass spectrometry data confirms the model built for the long loop extension of S6. **g**, The crosslinking network of S6 identified in the previous work[16], with the sequence range (1-167) resolved in the map colored in green.

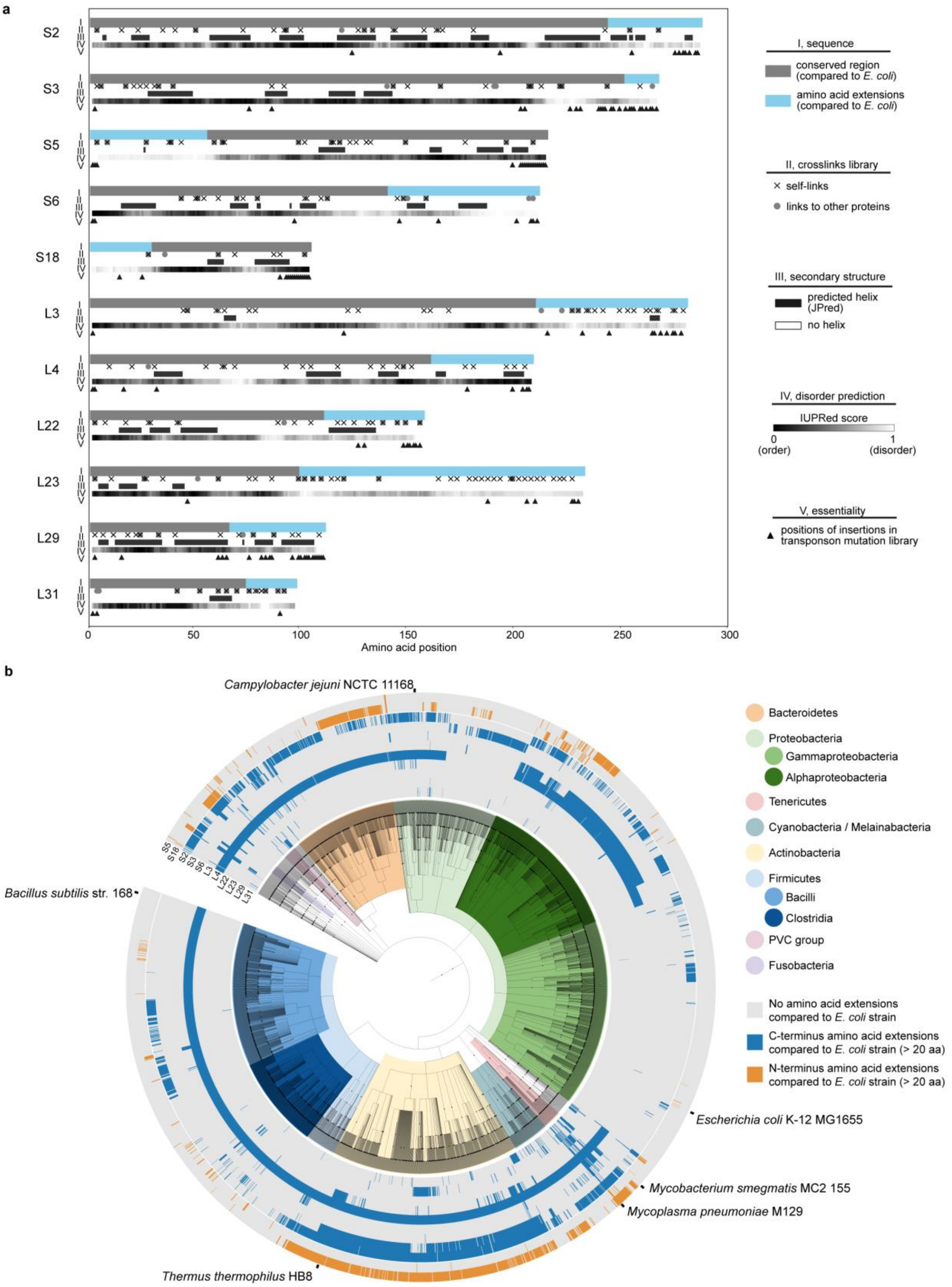


**Extended Data Fig. 3 | Ribosomal protein extensions.**

**a**, Eleven ribosomal proteins in *M. pneumoniae* have sequence extensions compared to *E. coli*. Along the sequences, crosslinks identified by in-cell crosslinking mass spectrometry, secondary structure prediction, disorder prediction, and sequence essentiality prediction based on a transposon mutation library are displayed (quantified in Supplementary Table 1). **b**, Extensions in the eleven ribosomal proteins are found throughout bacteria, but are not specific to any sub-groups.

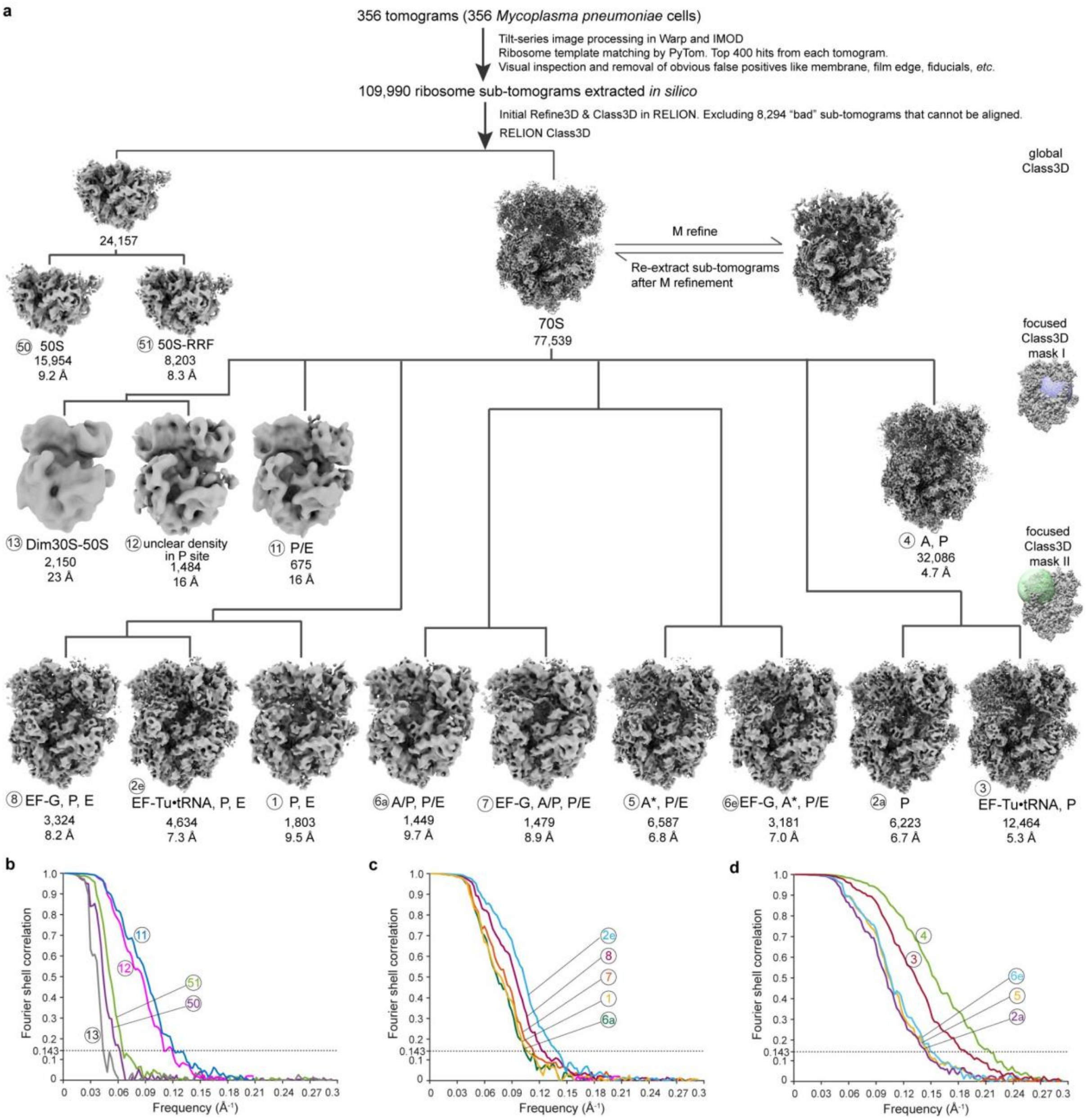


**Extended Data Fig. 4 | Classification and refinement of ribosomes in untreated cells.**

**a**, A diagram of the image processing workflow. The classification process can be grouped into three tiers: global classification, focused classification on the tRNA path region (mask I), and focused classification on the elongation factor and A/T tRNA binding region (mask II). For each class, the unique number identifier and class name are assigned for tracking. The particle number and the global map resolution at 0.143 FSC are provided for each class. **b-d**, FSC curves for all refined maps.

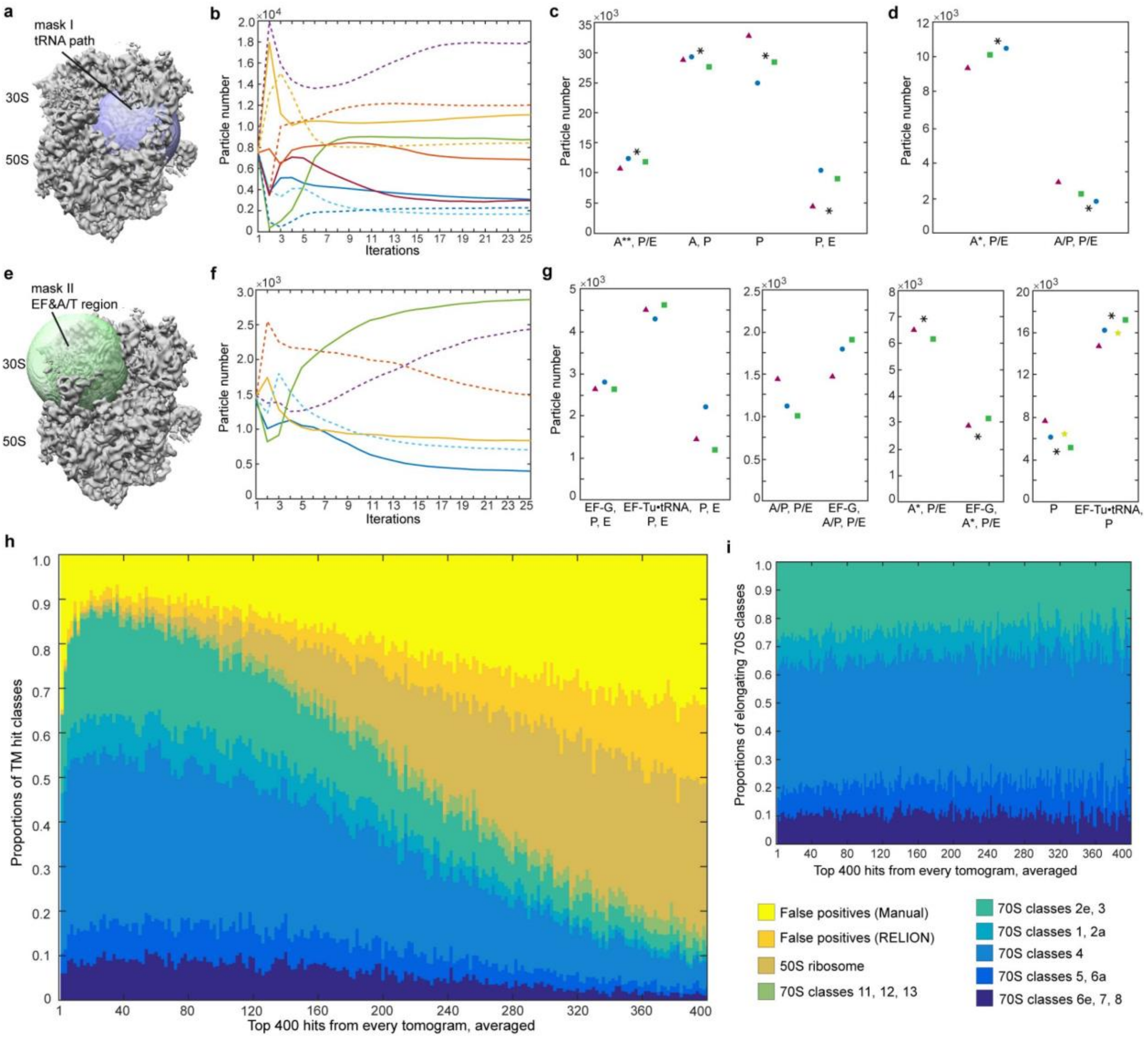


**Extended Data Fig. 5 | Validation of ribosome detection and classification.**

**a**, Mask I for focused classification on tRNA path region. **b**, A representative RELION classification job with mask I. Each line indicates the change in particle numbers in one class in 25 iterations. Classes that show the same structure were grouped according to the tRNA occupancy ("A**, P/E", "P, E", "P", "A, P"). **c**, Results of the classification job as shown in **b**, and of three additional parallel jobs. **d**, Results of following classification jobs that further classify the "A**, P/E" class into "A*, P/E" and "A/P, P/E". **e**, Mask II for focused classification on EF and A/T tRNA sites. **f**, Changes in particle numbers with iterations in a representative classification job with the mask II. **g**, Results of parallel focused classification jobs with mask II. **h**, Distribution of the ribosome classes against template matching cross-correlation scores employed for ribosome localization. For each of the 356 tomograms of untreated cells, 400 highest scoring hits were extracted and ranked. Obvious false positives were manually excluded first. Additional false positives were identified during RELION classification. 70S classes that are structurally similar were grouped in the plot for better visualization. **i**, Same as **h**, but only showing the 70S classes in the elongation phase. The proportions of different 70S classes remain stable across the top 400 hits, demonstrating that the classification results are not biased by the ribosome picking.

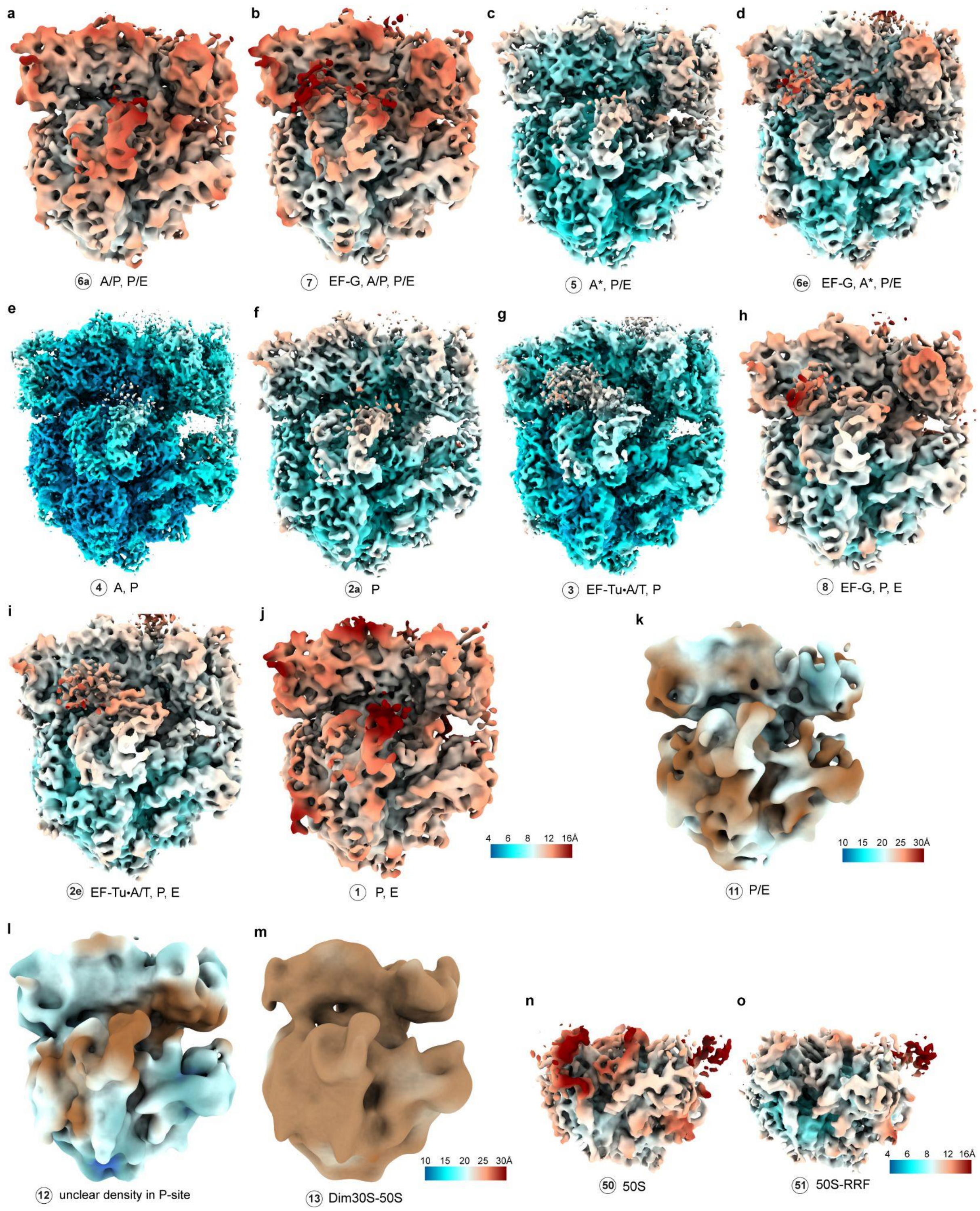


**Extended Data Fig. 6 | Local resolution maps for ribosome classes in untreated cells.**

Maps of the 15 classes determined in the untreated cellular dataset (Extended Data Fig. 4), colored by local resolution calculated in RELION.

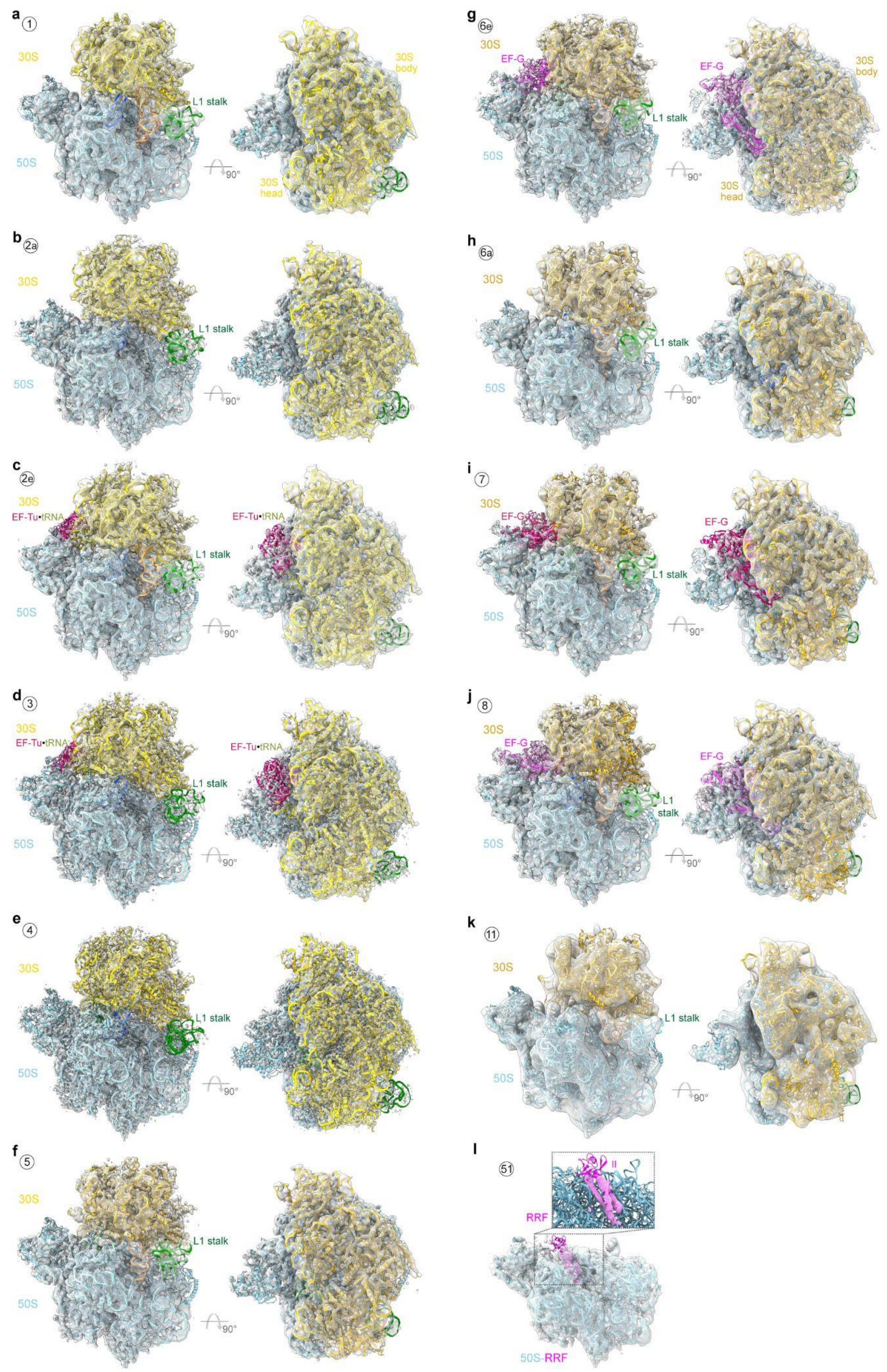


**Extended Data Fig. 7 | Model building for ribosome classes in untreated cells**

**a-j**, Models of the ten ribosome classes in the elongation phase by flexible fitting. **k**, The model for the ribosome with hybrid P/E-site tRNA. **l**, Free 50S in complex with ribosome recycling factor.

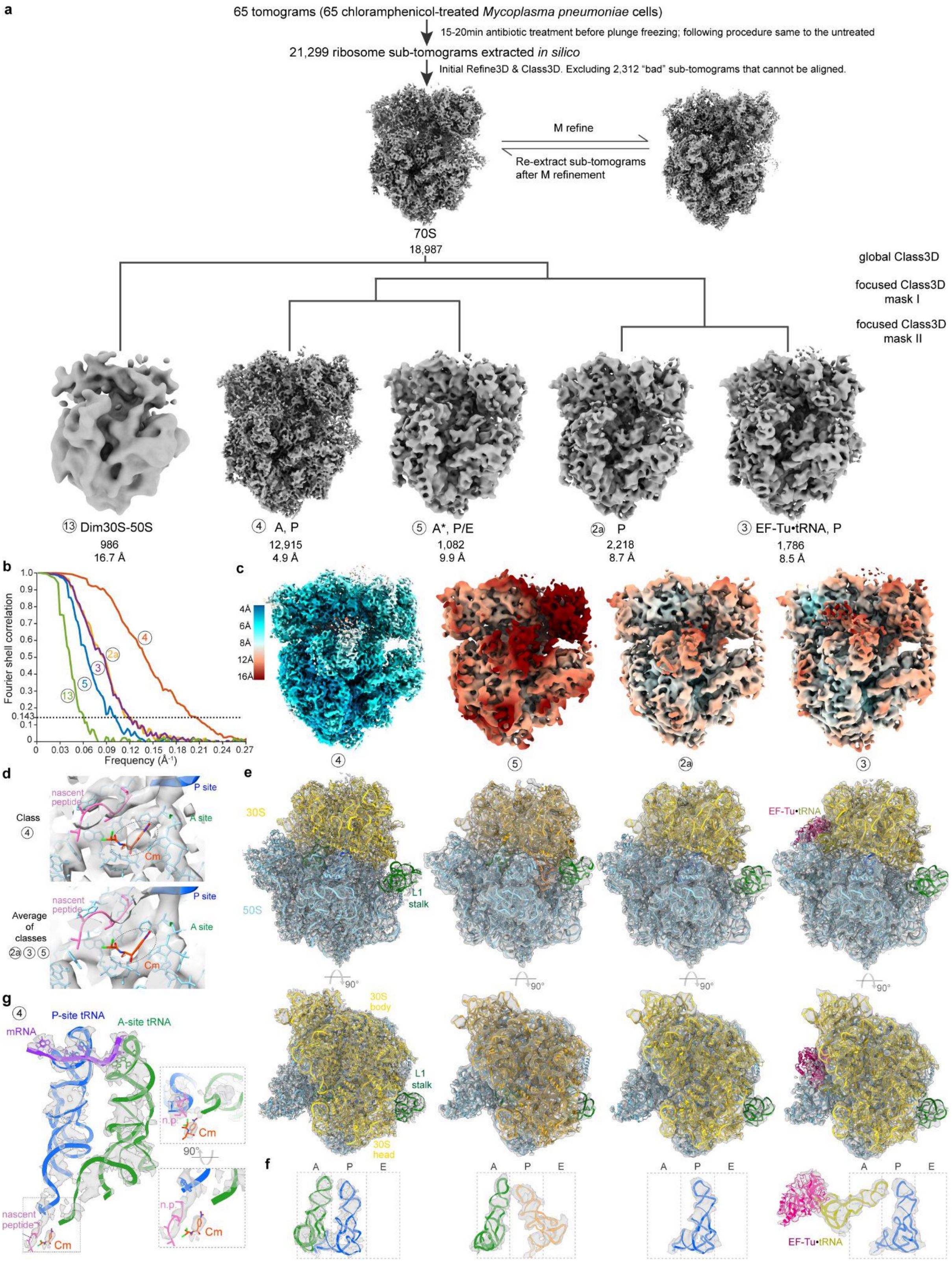


**Extended Data Fig. 8 | Classification, refinement and modeling of ribosomes in Cm-treated cells.**

**a**, The processing workflow of the Cm-treated dataset. The sub-tomogram classification and refinement are identical to those developed for the untreated dataset (Extended Data Fig. 4). **b**, FSC curves for all classes calculated following RELION refinements. **c**, Local resolution maps. **d**, Density of the Cm molecule is resolved in the major "A, P" class, but not in the other three minor classes. **e**,

Models built for the four classes, fitted into their corresponding densities. **f**, Elongation factors and tRNAs in the four classes. **g**, In the predominant "A, P" class, mRNA, A- and P-site tRNA, the nascent peptide chain, and the Cm drug are well-resolved. The nascent chain has strong continuous density linked to the P-site tRNA, resulting from the inhibition of peptidyl transfer by the Cm molecule.

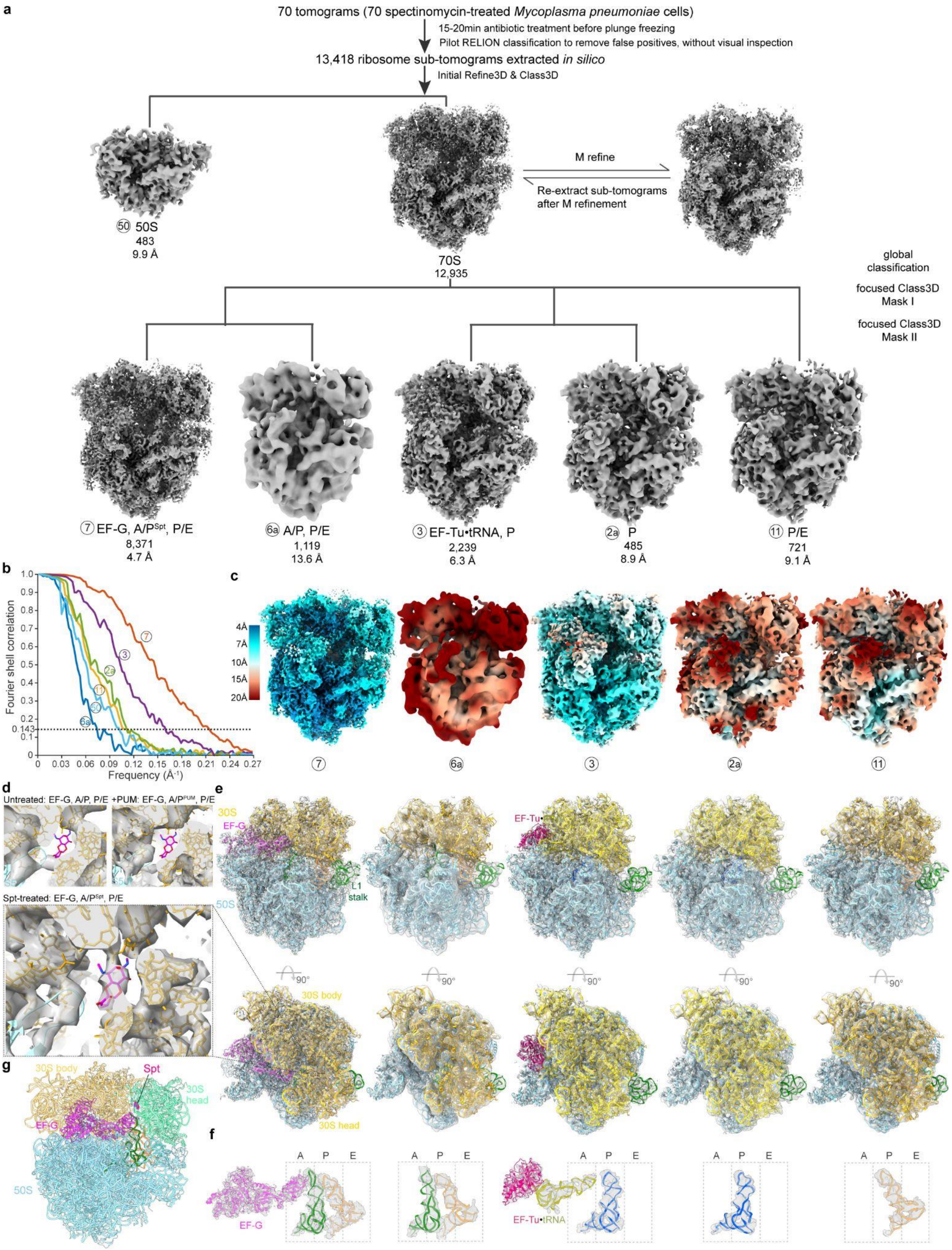


**Extended Data Fig. 9 | Classification, refinement and modeling of ribosomes in Spt-treated cells.**

**a**, The processing workflow for the Spt-treated dataset. The sub-tomogram classification and refinement are identical to those developed for the untreated dataset (Extended Data Fig. 4), excluding the manual inspection step after template matching. **b**, FSC curves for all classes calculated following refinements. **c**, Local resolution maps for Spt-treated 70S classes. **d**, Density corresponding to the Spt molecule (magenta) is clearly resolved in the major "EF-G, A/$P^{Spt}$, P/E" class. Density from the untreated and PUM-treated data, where Spt is not present, are shown for comparison. **e**, Models built for the five Spt-treated 70S classes. **f**, Elongation factors and tRNAs in the five classes. **g**, The model of "EF-G, A/$P^{Spt}$, P/E" shows Spt binds to the 30S neck region, confirming its role in inhibiting 30S head dynamics and mRNA translocation.

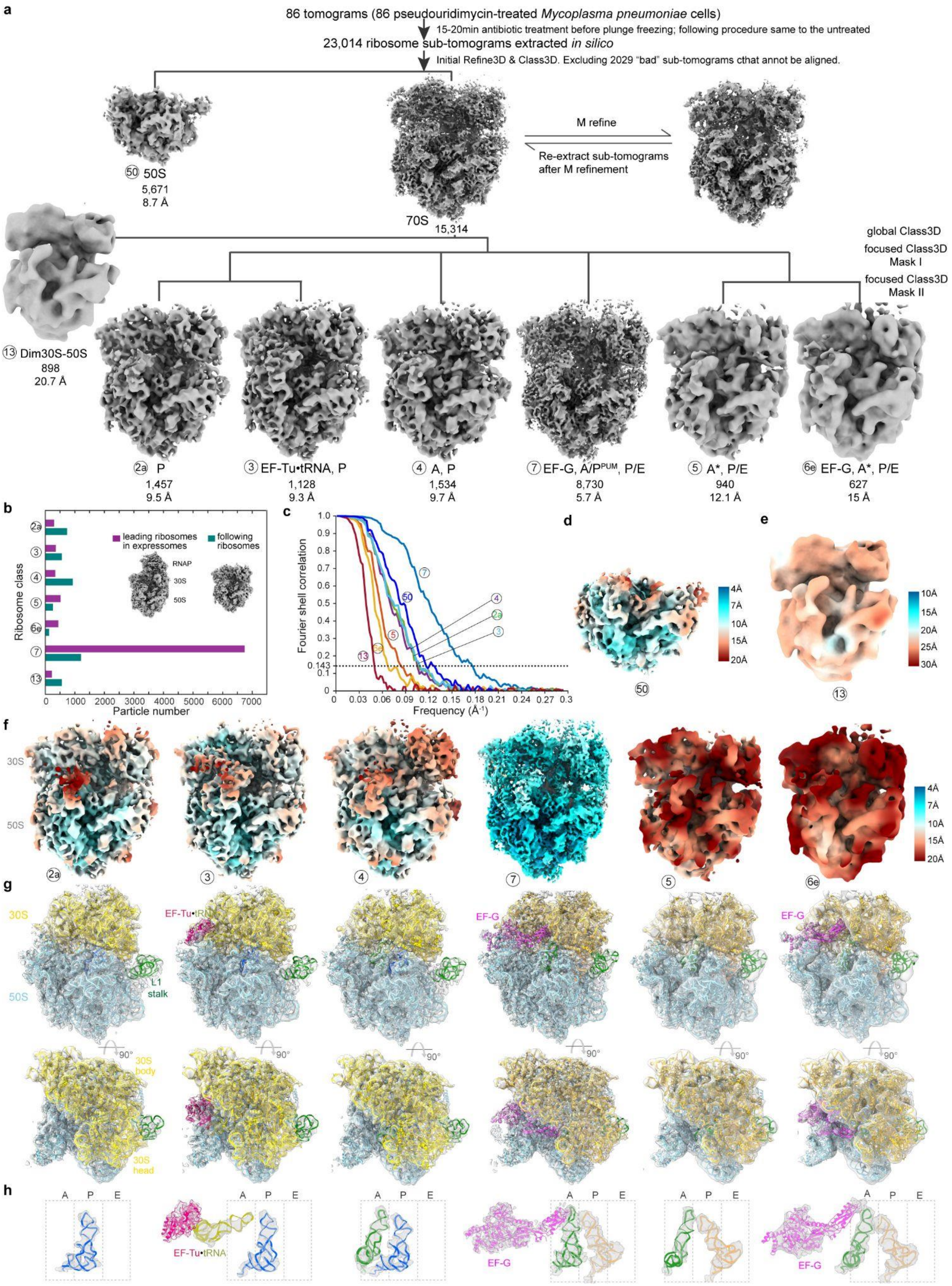


**Extended Data Fig. 10 | Classification, refinement and modeling of ribosomes in PUM-treated cells.**

**a**, Diagram of the processing workflow for the PUM-treated dataset. The sub-tomogram classification and refinement are identical to those developed for the untreated dataset (Extended Data Fig. 4). **b**,

Class distribution of elongation states for ribosomes collided with a stalled RNA polymerase (purple) and for the remaining ribosomes (green). **c**, FSC curves for all classes calculated following refinements. **d-f**, Local resolution maps. **g**, Models built for PUM-treated 70S classes. **h**, The elongation factors and tRNAs in the six 70S ribosome classes in PUM-treated cells.

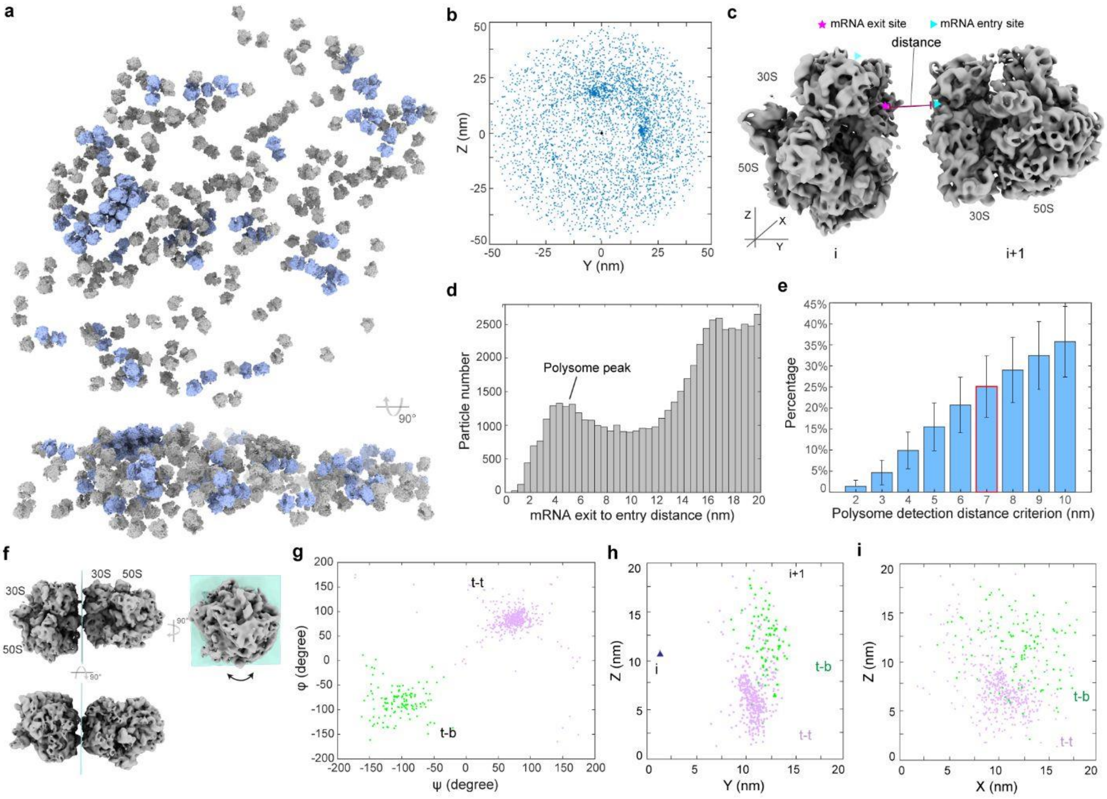


**Extended Data Fig. 11 | Spatial analysis of in-cell ribosomes and polysomes.**

**a**, Map of 70S ribosomes (grey) and detected polysomes (light blue) in a representative tomogram. **b**, Distribution of all neighboring ribosomes within 50 nm center-to-center distance. **c**, Illustration of the polysome detection criterion. The distance from the mRNA exit site of the preceding ribosome (i) to the mRNA entry site of the following ribosome (i+1) is calculated and used to define whether they belong to the same polysome. **d**, Histogram of distances from the mRNA exit site to the mRNA entry sites of all neighboring ribosomes. The peak at 4 nm is attributed to the formation of closely assembled polysomes. **e**, Percentages of ribosomes detected as polysomes using different distance thresholds. Mean and standard deviation across 356 untreated cells are shown. A threshold of 7 nm was selected to define polysomes. **f**, Orientations of the following ribosome relative to the preceding ribosome in polysomes. The major flexibility is the rotation (up to 360°) around a plane (light blue) perpendicular to the mRNA exit site of the preceding ribosome, which defines the "t-t" and "t-b" configurations in this study. **g**, Distribution of relative rotations of the following ribosome to the preceding ribosome in polysomes. The two clusters correspond to the defined "t-t" and "t-b" configurations. **h-i**, Positions of the following ribosomes (i+1, colored by different rotations) relative to the preceding ribosome (i, triangle indicates the mRNA exit site) in polysomes.

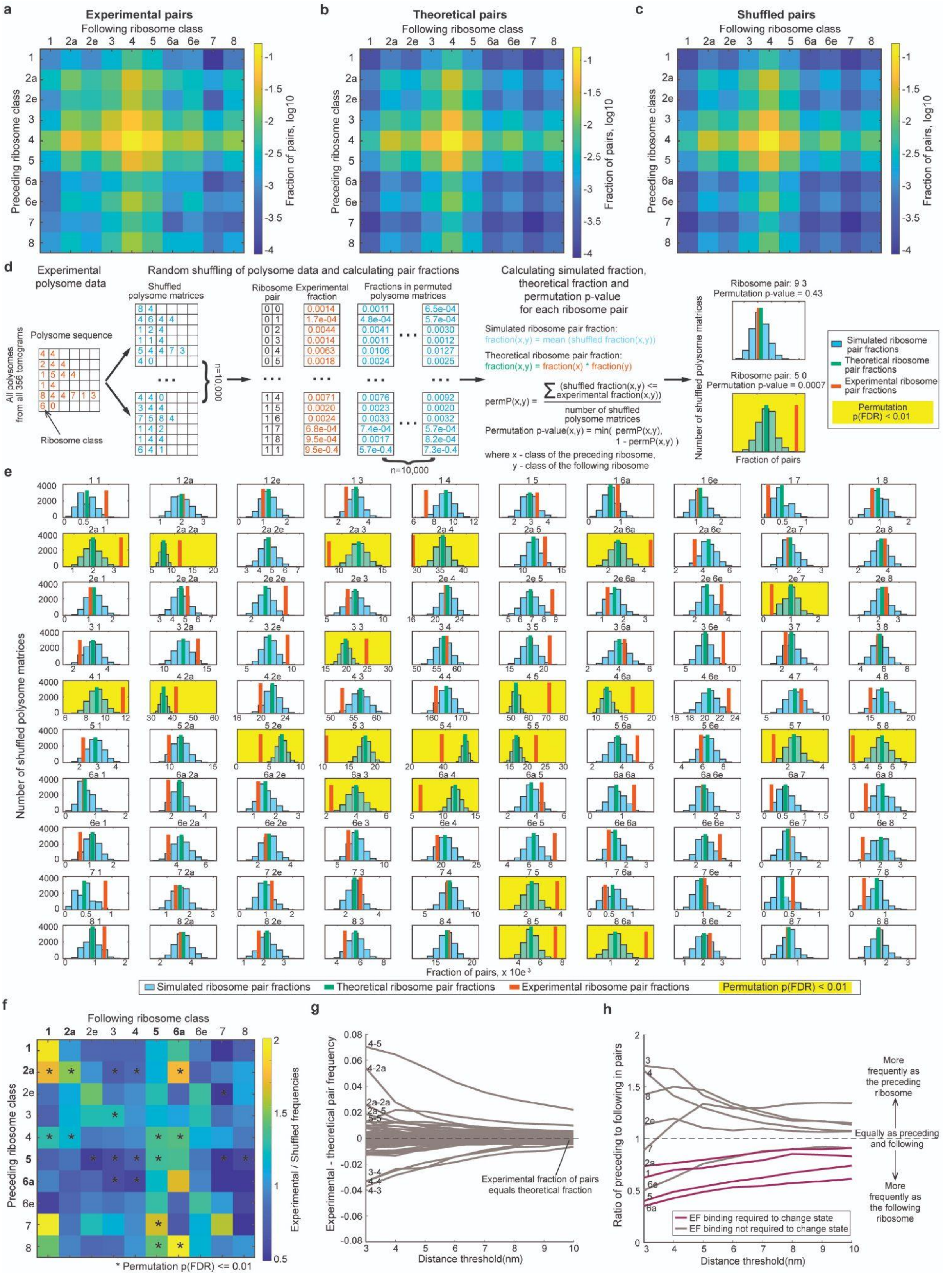

**Extended Data Fig. 12 | Correlation of translation elongation states within polysomes.**

**a**, Occurrence frequencies of elongation state pairs of two adjacent ribosomes (preceding ribosome i *vs.* following ribosome i+1) within polysomes calculated from the experimental data. **b**, Theoretical frequencies of elongation state pairs if there is no cross-influence within the polysome and ribosome state pairs form randomly. **c**, Frequencies of elongation state pairs calculated after random shuffling of the experimental data. **d,** Schematic representation of polysome shuffling analysis and permutation *p*-value calculation. Detailed procedure can be found in the Materials and Methods. **e**, Comparison of the experimental and shuffled pair fractions for all major pairs. **f**, Fold changes between the experimental and shuffled pair fractions from the shuffling experiments. Ribosomes of states that need elongation factor binding to proceed (states 1 and 2a for EF-Tu, and states 5 and 6a for EF-G) are more frequently engaged as the following ribosomes. **g**, Difference between experimental and theoretical pair frequencies when using different distance thresholds for polysome definition. **h**, Ratios of the number of ribosomes in each state being the preceding one against the following one in polysome pairs (across all polysomes), calculated using different distance thresholds to define polysomes. Symmetric engagement as the preceding and following ribosome results in a ratio of 1.

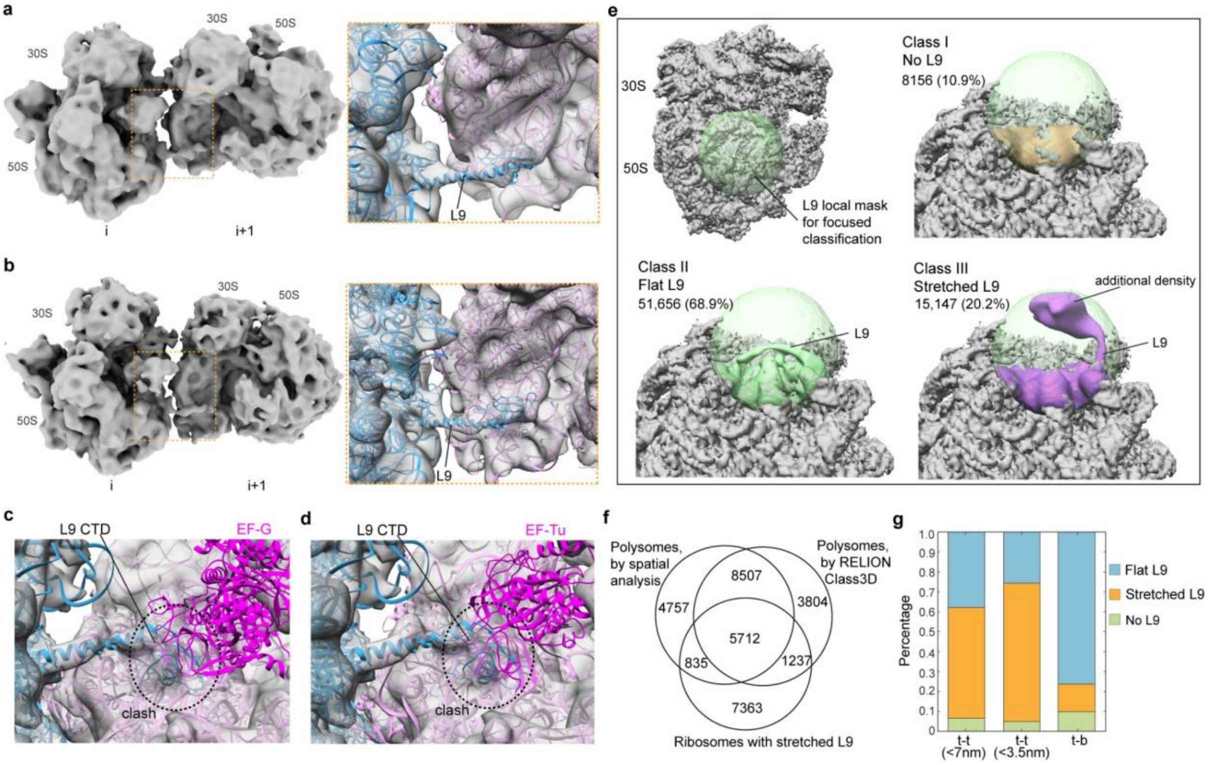


**Extended Data Fig. 13 | Stretched L9 within the ribosome-ribosome interface in polysomes.**

**a-b**, Two representative di-ribosome classes with a better resolved ribosome-ribosome interface. The stretched conformation of ribosomal protein L9 of the preceding ribosome is highlighted. **c**, Steric clash between the C-terminal domain of the stretched L9 and the superposed EF-G according to the "EF-G, A/P, P/E" structure. **d**, Clash between L9 and EF-Tu as in "EF-Tu•tRNA, P" class. **e**, *De novo* focused classification on the L9 region using a local spherical mask (light green) of all 70S ribosomes from untreated cells. Densities from the three resulting classes are displayed on the ribosome surface: no L9 resolved (I), flat L9 (II) and stretched L9 (III). The additional density in contact with the stretched L9 stems from the neighboring ribosome. **f,** Overlap between ribosomes within polysomes independently defined based on spatial analysis (Extended Data Fig. 11), polysomes from *de novo* RELION classification, and ribosomes classified with a stretched L9. **g**, Distribution of different L9 conformations in adjacent ribosome pairs in polysomes with the "t-t" and "t-b" configurations. The stretched conformation is more frequently observed in the "t-t" ribosome pairs, especially for more compacted pairs with less than 3.5 nm mRNA exit-to-entry distance.

#### Extended Data Movies

Extended Data Movies are provided as separate files.

**Movie 1 | Visualizing the translation elongation cycle in untreated cells.**

Maps of ribosome classes within the elongation phase are displayed in a sequential manner to illustrate the continuous structural changes during translation elongation. Maps are low-pass filtered to 10 Å for normalization.

**Movie 2 | Structural dynamics of translation elongation in untreated cells.**

The atomic models for ribosome intermediates within the elongation phase in untreated cells are displayed sequentially to illustrate the structural dynamics during the translation elongation cycle.

#### Extended Data Tables

Extended Data Tables are provided in a separate Excel file.

Extended Data Table 1 | Cryo-EM data collection, refinement and validation statistics of the high-resolution ribosome averages.

Extended Data Table 2 | Cryo-EM data collection, refinement and validation statistics of ribosome classes in untreated cells.

Extended Data Table 3 | Cryo-EM data collection, refinement and validation statistics of ribosome classes in chloramphenicol-treated cells.

Extended Data Table 4 | Cryo-EM data collection, refinement and validation statistics of ribosome classes in spectinomycin-treated cells.

Extended Data Table 5 | Cryo-EM data collection, refinement and validation statistics of ribosome classes in pseudouridimycin-treated cells.
